# Stage-specific habitat use of the Mountain Plover in Colorado, USA

**DOI:** 10.64898/2026.01.29.702628

**Authors:** Casey M. Weissburg, Allison K. Pierce, Michael B. Wunder, Claire W. Varian-Ramos

## Abstract

Habitat structure, food availability, and predation risk have spatiotemporal variation on the landscape that creates tradeoffs between risk and rewards for animals as they use habitat. These tradeoffs are associated with survival consequences and can vary by life or breeding stage, but stage-specific shifts in these relationships are often not considered in studies of habitat use and survival. We investigated habitat use for nests and broods in the Mountain plover as an example of a ground-nesting species with precocial, mobile young to explore breeding stage-specific responses to vegetation structure, food availability, and predation risk. We located and monitored nests and broods at two study sites in Colorado occupied by geographically separated breeding populations of plovers. We quantified the three covariate categories across standardized site-wide grids in 2021 and 2022. We employed a resource selection analysis to evaluate 10 *a priori* working hypotheses for how environmental covariates may influence the habitat used by plovers for nesting and brood-rearing. Model comparison results suggest that habitat use relative to availability is best explained by a linear relationship with vegetation structure while conditioning on breeding stage and site, with no effect of food availability or predation risk detected in our dataset. Specifically, probability of use for both nest and brood sites was highest in areas with taller vegetation, contrary to previous research about nest site selection. These unexpected results may be due to the unbalanced sample sizes of the two sites, as previous separate studies have shown opposing habitat use strategies, or due to the coarseness of our covariate data. Regardless of limitations, these results emphasize the importance of investigating stage-specific and potentially site-specific shifts in habitat use patterns for species with precocial young.

## INTRODUCTION

Understanding the environmental factors that underlie species distributions can facilitate identification of critical habitat and support ecological forecasting. Habitat use is a complex process, however. An animal must manage balances of energy, heat and water, and risk to grow, survive and reproduce (Cody, 1981; Fletcher & Fortin, 2018; Piersma, 2012). Numerous factors can influence an individual’s ability to balance needs and dangers, including vegetation structure (Clark & Shutler, 1999; Cody, 1981; Martin, 1993; Piersma, 2012), food availability (Cody, 1981; Hutto, 1977; Swift et al., 2023), and predation risk (Creswell & Quinn, 2013; Mönkkönen et al., 2007; Swift et al., 2018).

Predation risk creates tradeoffs for animals between managing energy and water balances and avoiding danger, as spatiotemporal variation creates a dynamic landscape of fear (Laundré et al., 2010; Piersma, 2012). For example, some birds use nest sites at an optimal distance from a predator to gain incidental nest protection from egg predators (Swift et al., 2018; Wilde et al., 2022) sometimes even at risk to incubating adults (Mönkkönen et al., 2007). Others use habitat with vegetation features that offer crypsis to minimize risk during incubation or brood-rearing (Colwell et al., 2005; Hood & Dinsmore, 2007; Knopf & Miller, 1994; Skrade & Dinsmore, 2013; Winton et al., 2000; Webber et al., 2013). Others still use habitat with higher food availability regardless of risk or apparent lack of fitness benefits (Bloom et al., 2013; Chalfoun & Martin, 2007), while some balance risk and reward by using habitat with lower risk or disturbance while optimizing availability of foraging habitat and success (Creswell & Quinn, 2013; Swift et al., 2023).

The balance between risk and reward can also vary by life stage (Robinson et al., 2021), breeding stage (Paasivaara & Pöysä, 2008; Webber et al., 2013), or even time of day (Nugent et al., 2022). Species with precocial young may attempt to move mobile broods near foraging areas with high food availability (Cohen et al., 2009; McIntyre & Heath, 2011), away from areas with higher disturbance (Webber et al., 2013), or away from previously protective nesting associations that endanger vulnerable chicks (Swift et al., 2018; Wilde et al., 2022). These tradeoffs influence individual survival and fitness, and in turn population growth (Bloom et al., 2013; Chalfoun & Martin, 2007; Clark & Shutler, 1999; Devries et al., 2018; Martin, 1998).

Yet investigations that aim to better inform management and conservation of habitat for declining species often do not consider individual responses to predation risk, comprehensive evaluation of responses to both risks and rewards, or potential life stage-specific habitat requirements. Our study therefore uses a comprehensive approach to evaluate multiple working hypotheses about three features of the landscape that influence habitat use: vegetation structure, potential food availability, and predation risk, while evaluating the strength of evidence for a dependence on breeding stage.

We studied the Mountain plover (*Anarhynchus montanus*), a ground-nesting shorebird species with precocial, mobile young (Knopf & Wunder, 2023) with an unexplored potential for breeding stage-related shifts in habitat use. Previous research across the known geographic range has shown a consistent use of habitat with a high proportion of bare ground and short, sparse vegetation during the nesting period (Andres & Stone, 2010; Augustine & Derner, 2012; Duchardt et al., 2020; Graul, 1975; Knopf & Miller, 1994; Manning & White, 2001) and wintering period (Wunder & Knopf, 2003; Lyons et al., 2025), but there remains no research on how this species responds to predation risk (Parker et al., 2019) and very little research during the brood-rearing stage. Studies of brood movements are limited (Dreitz et al., 2005; Knopf & Rupert, 1996), and just two studies have examined habitat use relative to availability during this stage (Schneider et al., 2006; Sordahl, 1991), with conflicting results.

Investigation into stage-specific habitat use is of particular interest for the Mountain plover. The species is estimated to have declined as much as 70% since the 1970s (Andres & Stone, 2010; Knopf, 1994; Partners in Flight, 2019; Rosenberg et al., 2019). Population growth in the plover is most elastic to adult survival during migration and chick survival (Dinsmore et al., 2010), yet research and management strategies have remained focused on nest survival, which has little influence. While some studies have estimated baseline chick survival rates (Dreitz, 2009; Knopf & Rupert, 1996; Lukacs et al., 2004; Miller & Knopf, 1993; Ruff, 2016), little is known about how plovers respond to spatial variation of habitat features during the brood-rearing stage.

Our research therefore investigated potential differences in habitat use, and the relative influence of vegetation structure, food availability, and predation risk, during stages that differ in mobility, from stationary (nests) to relatively limited in movement with respect to the study area (non-flighted but mobile precocial chicks). We located and monitored nests and broods for two consecutive years (2021-2022) at two study sites in Colorado occupied by geographically separated breeding populations of plovers. We used a resource selection framework in a series of ten *a priori* models to evaluate the relative strength of evidence for which habitat features best predict plover habitat use relative to what was available on the landscape, and the degree to which the effects of these habitat features depended on breeding stage.

## MATERIALS & METHODS

### Study areas

We collected data at two study sites in east-central Colorado, where greater than 60% of known breeding populations occur (Andres & Stone, 2010; Figure 1). Sites represent similar but distinct grassland habitats, the eastern plains and high-elevation intermountain basins. The eastern plains study site, Chico Basin Ranch (CHBR), encompasses ∼15 sq km (38°28’43”-32’2” N, 104°23’16”-25’16” W) of shortgrass prairie characterized by black-tailed prairie-dog (*Cynomys ludovicianus*) colonies in a grazed mosaic of bare ground, short grasses (*Bouteloua gracilis, Aristida purpurea*), and sparse shrubs and cacti. The high plains study site, South Park (SP), is on public land at James Mark Jones State Wildlife Area and adjacent Bureau of Land Management (BLM) pastures with an area of ∼25 sq km (39°7’15”-12’28” N, 105°48’41”-52’5” W). This intermountain basin sits at an elevation of 2,800 meters and is characterized by short grasses (*Muhlenbergia filiculmis*, *M. montana, B. gracilis*), extensive stands of very short forbs and shrubs (*Artemisia frigida*, *Chrysothamnus viscidiflorus*), and bare ground that is relatively more limited to dry alkaline playa lakes, flood washes and ungulate wallows (Wunder et al., 2003).

**Figure 1.**
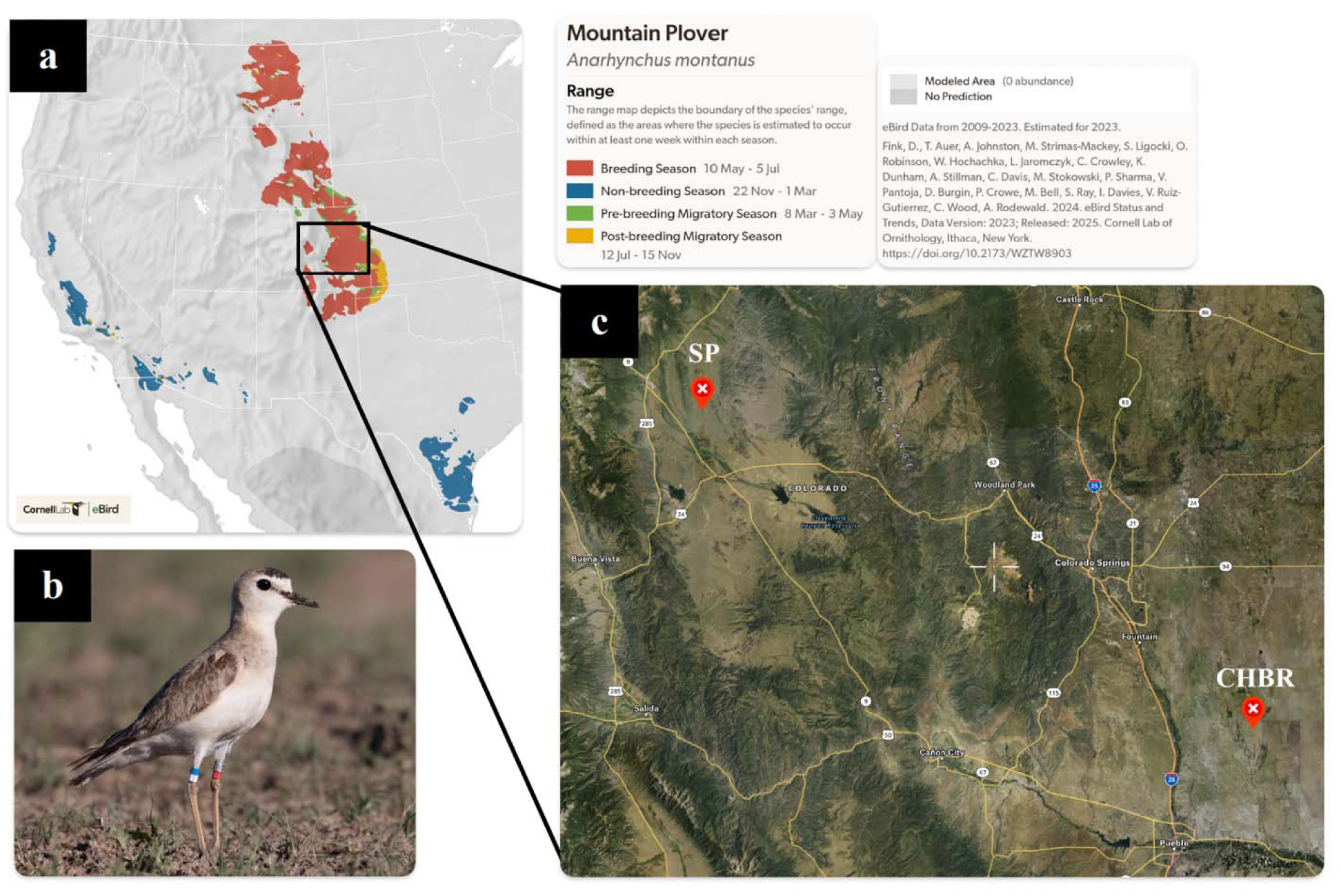
Geographic arrangement of study sites. Visualization of (a) the species range (Fink et al. 2023) of the (b) Mountain plover, with (c) an inset of central Colorado displaying the geographic arrangement of our study sites (CHBR and SP).

### Nest searching & monitoring

Nest searching and monitoring efforts followed the methods of Dinsmore et al. (2002). Nesting phenology differs between study sites; we conducted systematic searches for nests during April-June (CHBR) and late May-July (SP) with some overlap, searching all available habitat within study site boundaries and repeating surveys once per week or every other week to account for low detection probability of this species. We recorded GPS locations upon nest discovery and floated eggs in a column of water to estimate hatch date (Dinsmore et al., 2002). We checked nests every 3-7 days until they failed or hatched. We marked incubating adults with an aluminum USGS leg band and unique color combination for ongoing mark-recapture efforts.

### Brood monitoring

The highly mobile young of this species leave the nest as soon as they are dry (Graul, 1975), and adults typically move away from chicks when observers approach (Whittingham et al., 1999; CW, personal obs.), so we used radiotelemetry to evaluate habitat use directly from chick locations within the scope of the study area. We monitored broods for as many known nests as possible that hatched. We captured chicks by hand, most often from the nest cup to minimize disturbance and improve accuracy of aging. Broods discovered away from nests were also captured to be monitored, with chick age estimated by a combination of body mass, tibiotarsus length and personal experience with chicks of known ages (CW, in process). We marked all captured chicks with an aluminum USGS leg band; we deployed unique color combinations only within one week of fledging so as not to increase visibility to predators.

Plover broods move together as a unit (Graul, 1975); we thus selected one chick from each brood for deployment of a 0.48 g VHF radio-transmitting tag (Lotek Ag379 PicoPip, ∼42-day battery life). We affixed tags by application of superglue to a small patch of skin on the back clipped free of down, as in previous studies of shorebird chicks (Miller & Knopf, 1993; Whittingham et al., 1999). We tracked tagged broods via radio telemetry until we confirmed fledging (sustained flight for at least 10 meters) or death, obtaining approximately one fix per day whenever logistically possible. We maintained a minimum distance of 100 m when possible, to minimize pushing broods (Sordahl, 1991; Whittingham et al., 1999; CW, personal obs.) and approached only if we suspected mortality. If a tagged chick was lost, we attempted to relocate the attending adult to tag a sibling if any remained, since our sampling unit was the brood, not the individual chick.

### Habitat feature indices

The purpose of this study was to evaluate the relative influence on habitat use of spatial variation in three categories of habitat features: predation risk, food availability, and vegetation structure. See Figure 2 for a heuristic of the sampling design. To characterize the spatial variation of covariates of interest, we quantified all of them across a standardized 600-by-600-meter grid of sampling locations (CHBR, n = 38 locations; SP, n = 70 in 2021, 80 in 2022 when site boundaries were expanded). Following field collection, we rasterized covariate data across the grids and estimated a finer resolution of 300-by-300 meters using inverse distance weighted interpolation in QGIS (See Supporting Information Figures S1-4 for site-specific and year-specific rasters). We established this resolution for the sampling grid to reflect the scale of daily movements of plover broods estimated in previous works (∼300 m per day; c.f. Dreitz et al., 2005; Graul, 1975; Knopf & Rupert, 1996) and we defined the boundaries of the grid by ecological shifts to habitat unsuitable for plovers (forested ridges at SP; uninterrupted tall grass and dense shrubs at CHBR) as described in previous research (Andres & Stone, 2010; Augustine & Derner, 2012; Duchardt et al., 2020; Graul, 1975; Knopf & Miller, 1994; Manning & White, 2001). Due to logistical constraints, collection could not span the duration of the breeding season and thus temporal variation of covariates and habitat use responses cannot be assessed by this study.

**Figure 2.**
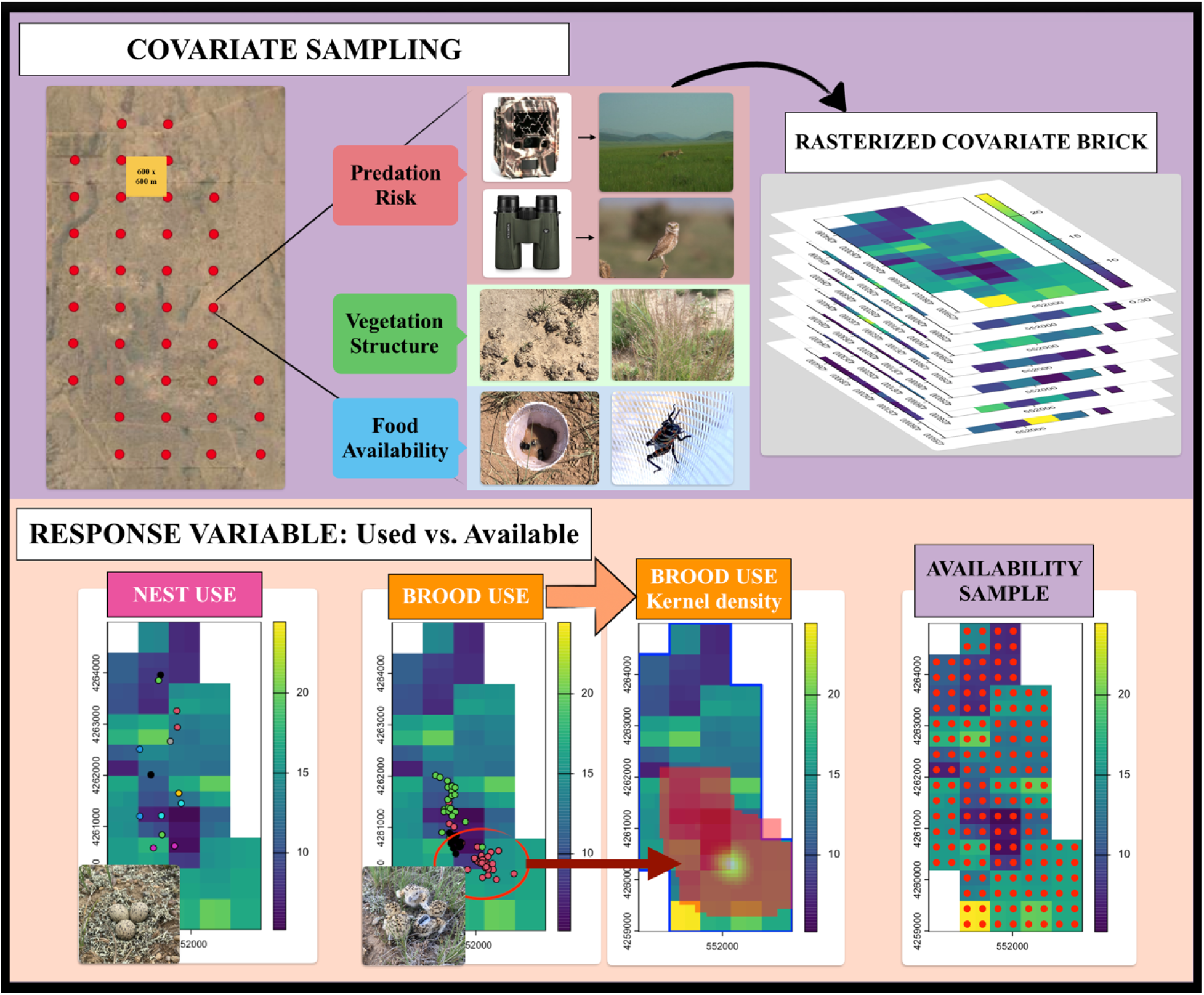
Heuristic of sampling design. A heuristic diagram of the sampling design for variables, using CHBR in 2021 as an example. We sampled all covariates at every location on a standardized site-wide grid and then rasterized all seven into a raster brick (top panel). The response variable (bottom panel) consisted of nest and brood locations used by plovers contrasted with availability samples extracted from every cell of the raster brick.

### Predation risk

Plover nests and chicks have numerous predators, both mammalian and avian. We thus characterized spatial variation of predation risk with indices for both mammalian and avian predators. We quantified indices of risk as encounter rates, because we were interested in the influence of *perceived* risk, as opposed to predator density. We kept indices separate since the data collection methods were different and additionally because the detection by plovers of mammalian and avian predators could differ in unquantifiable ways.

We quantified the **mammalian predator risk index** by passive detection, using a single infrared trail camera deployed at each sampling location on the grid. Cameras had a viewshed of 30 m and were affixed north-facing and approx. 2 ft above ground to a T-post. Capture periods ran continuously for three to four weeks, starting the week before or of the first nest hatching (CHBR, mid-May; SP, mid-June); active sampling in SP had to be rotated due to limited camera and battery supply. The duration of active periods varied from camera to camera due to interference from domestic and wild animals; therefore, we averaged mammalian predator detections across active trap days to obtain a single index value of mammalian predator detections per trap day at each sampling location, pooling all species detected (coyote, badger, swift fox, bear).

We quantified the **avian predator risk index** from standardized 10-minute point count surveys, conducted at all sampling locations once per week for five to six weeks starting the week before or of the first nest hatching, between the hours of 0900 and sunset and not in rain or winds >25mph. Double-counting within surveys was avoided; however, we included repeated observations of individual birds between surveys, as a measure of perceived risk. We pooled avian predator detections across all species (*Buteo* hawks, eagles, Northern harrier, all falcons, Common raven, Burrowing owl, vultures) and adjusted detections by the number of surveys conducted (five or six) to obtain average detections per survey for each location on the grid.

### Food availability

Mountain plovers are generalist insectivores of surface-dwelling arthropods (Stoner 1941; Baldwin 1971; Olson 1985; Knopf 1998), so to characterize spatial variation of food availability, we quantified two indices: 1) **dry biomass of all arthropods**, collected at every sampling location using three simple pitfall traps set 5 meters apart for a 72-hour collection period; and 2) **dry biomass of grasshoppers**, collected by net-sweeping along a 12-meter transect centered on each pitfall line. Grasshopper biomass was included separately since pitfall traps are not an effective sampling method for orthopterans (Schneider et al., 2006) and grasshoppers are known to be a key food item for this species (Olson, 1985). Due to logistical restraints, insect collection was conducted once each summer, during peak brood activity for each study site (mid-June at CHBR, mid-July at SP). We froze samples at collection and later excluded individual specimens with body lengths larger than 20 mm, according to prey size selection of similar-sized shorebird species (Lifjeld, 1984). We then dried sorted samples at 32 °C for 72 hours and weighed them to the nearest 0.01 g. We adjusted the mass of pitfall samples by the number of pitfalls successfully collected at the location, since some locations lost one or two of the three traps due to interference from animals or weather. This adjustment resulted in a single dry grams per effort value for each location on the grid.

### Vegetation structure

To characterize spatial variation of vegetation structure, we quantified three covariates: 1) proportion bare ground, 2) groundcover height, and 3) shrub density. Vegetation measurements were collected at the same time as insect sampling, during peak brood activity. We used a 0.25 m^2^ quadrant to collect four measurements of **proportion bare ground** and **groundcover height** at each sampling location, 5 meters from the central point in the four cardinal directions. Proportion bare ground was measured as the proportion (0-1) of ground within the quadrant that was bare of vegetation. Groundcover height was estimated as the average of three height (cm) measurements within each quadrant, of grass and forbs as well as shrubs that were <20 cm in height. We then averaged the four quadrant measurements to obtain mean proportion bare ground and groundcover height for each sampling location. We quantified **shrub density** at each sampling location by counting all shrubs (rabbitbrush, cholla, sagebrush, winterfat) that were 20 cm or taller within a 5-meter radius of the center (shrubs m^-2^). The 20 cm height cutoff for shrubs reflects the body length of the adult Mountain plover, such that our density of shrubs can be considered biologically relevant as visual obstruction between plovers and terrestrial predators.

While our indices of predation risk, food availability, and vegetation structure cannot describe temporal variation across the breeding season, they characterize the spatial variation of the environment available to plovers during the nesting and brood-rearing stages at a coarse resolution that matches average daily use patterns from previous research. We then extracted covariate data for all nest and brood coordinates from the year-specific, site-specific rasters.

### Statistical analysis

Data processing and analysis were performed within the R programming environment (v.4.3.2; R Development Core Team, 2023).

We implemented a use vs. available framework in a logistic regression (use = 1, available = 0) to model a *relative* measure of habitat use proportional to the probability of use (Johnson et al., 2006), with the aim to contrast plover use to what was available at known breeding sites (“second-order selection,” c.f. Fletcher & Fortin, 2018). Since our research question focused on the site-level use of habitat features where plovers are known to breed, and the scope and resolution of covariate data was relatively coarse, we characterized the “availability” in a systematic (non-random) manner at a spatial interval that matched the resolution of our covariate rasters, rather than generating arbitrary numbers of random locations until parameter coefficients stabilize (Northrup et al., 2013; Northrup et al., 2021). In this way, we censused habitat feature availability at every raster pixel, describing the entire availability distribution as estimated by our field methods (Benson, 2013; Fieberg et al., 2021; Street et al., 2021; Figure 2, bottom panel). Availability samples were extracted from year-specific rasters (CHBR, n = 152 both years; SP, n = 280 [2021] and 320 [2022]) and then duplicated and labeled as “Nest” or “Brood” to match the used data (CHBR, n = 304 per stage; SP, n = 600 per stage).

Plover use was represented by nest and brood locations. We had repeated observations for each brood, introducing pseudoreplication, as well as unbalanced sampling since the number of locations obtained from broods depended on duration of monitoring. Logistic regression assumes independence between data points, so we first intended to include a random intercept for nest/brood identification to improve model fit (Gillies et al., 2006). However, we encountered model singularity with even simple random intercepts, possibly due to the small sample size of CHBR or because the coarse resolution of the rasters meant that multiple brood locations had the same values.

As an alternative and novel solution to these challenges, we instead generated weighted averages for each sampling unit (a brood). We did this by leveraging the kernel density weights from the cells of kernel density estimated home ranges, to weight the covariate values extracted from those cells. Kernel density estimators use probability distributions to smooth the expected use of an animal (or in this case, a brood) around known use locations. Kernel density estimation of home range size does also assume independence of point locations (Worton 1989), which the fine temporal resolution of our brood data (near daily) is unlikely to satisfy (Swihart and Slade 1985); however, our use of the method here is to obtain *density of use*, represented by the kernel cell weights, rather than home range size. We estimated home ranges for each brood using the function *kernelUD()* in R package *adehabitatHR* with the default bandwidth estimator, and then paired the density of use weights with covariate values extracted from each cell in the home range to estimate kernel-density-weighted averages for each brood sampling unit. While this method results in some data loss through aggregation, we bypassed the problems of pseudoreplication and unbalanced sampling even with small sample sizes.

We collated nest covariate values and brood KD-weighted averages across years and sites into one dataset, along with the availability samples, so we could investigate effects of breeding stage. We standardized all covariate data to a standard normal distribution using the mean and standard deviation so that effect sizes could be compared among variables. We tested all covariate data of used and available units for correlation (Spearman’s rank correlation test) prior to modeling and we detected no significant correlations (Supporting Information: Figure S5).

We developed a series of *a priori* models representing the hypotheses that Mountain plover habitat use during the breeding season is best predicted by spatial variation of 1) predation risk; 2) food availability; 3) vegetation structure (linearly); 4) vegetation structure (quadratically); 5-8) each of the previous four relationships are dependent on an interaction with breeding stage (Nest vs. Brood). We did not include a year effect since the study only spanned two years and the term would only capture variation not explained by other, more biologically relevant terms. All models included an additive term for Site to control for geographic variation; Site could not be included as an interactive term in any models due to a large imbalance of samples (See Results: Data summary). We included an Intercept-only model as a null hypothesis and a global model containing all covariates interacting with Stage in the model set for comparison with our *a priori* hypotheses (see Table 1 for a complete list of hypothesis models and model terms included). We examined competitiveness of hypotheses using model comparison by Akaike’s information criterion for small samples (AICc; Anderson, 2008) to evaluate which hypothesis had the most strength of evidence to explain habitat use relative to spatial variation available in our dataset. We considered all models ≤2 ΔAICc competitive.

**Table 1.**
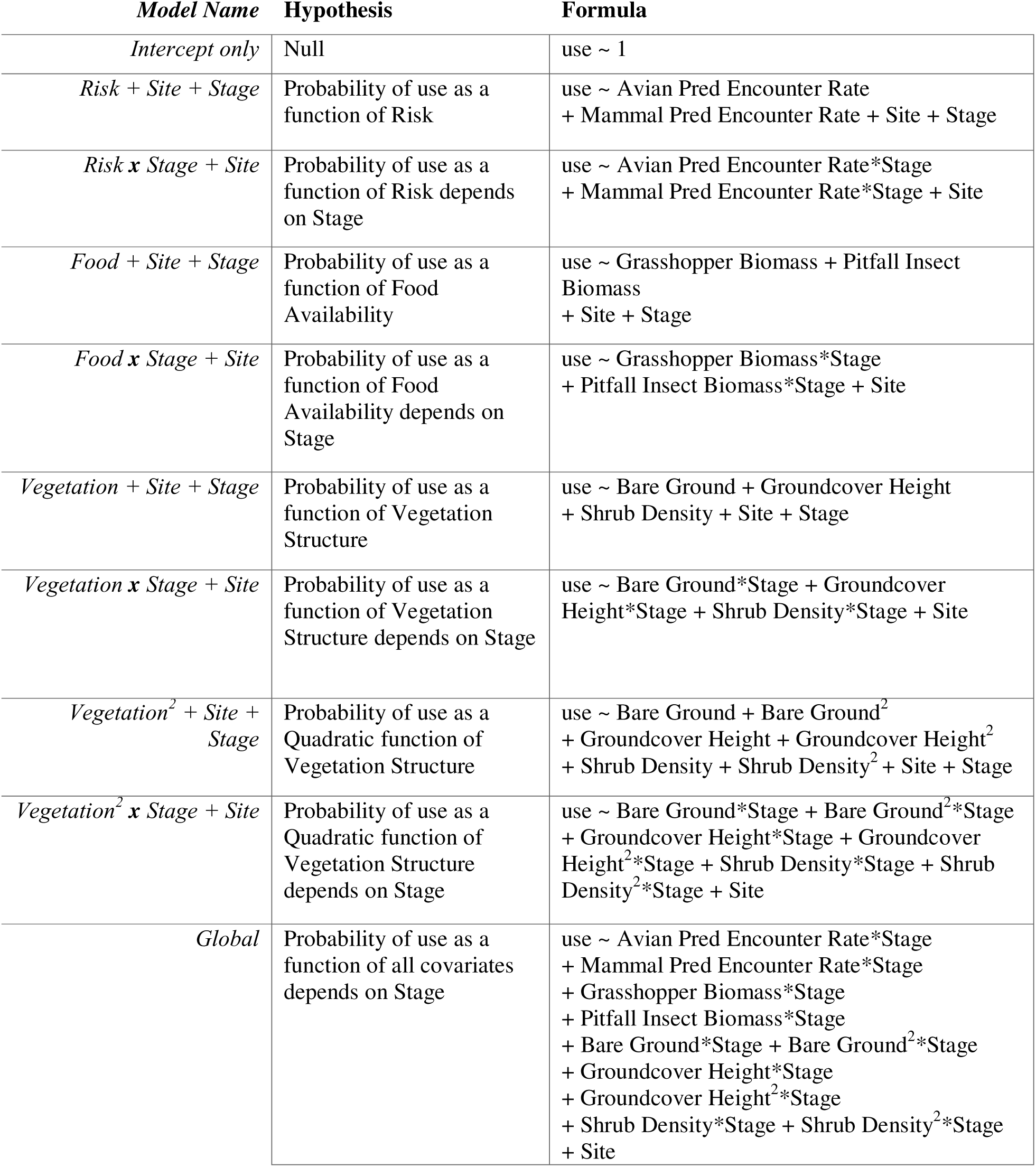
Full *a priori* model set. The full set of models considered for model comparison, listing specific hypotheses, short model name, and all terms included.

| *Model Name* | **Hypothesis** | **Formula** |
| --- | --- | --- |
| *Intercept only* | Null | use ~ 1 |
| *Risk + Site + Stage* | Probability of use as a function of Risk | use ~ Avian Pred Encounter Rate<br>+ Mammal Pred Encounter Rate + Site + Stage |
| *Risk x Stage + Site* | Probability of use as a function of Risk depends on Stage | use ~ Avian Pred Encounter Rate*Stage<br>+ Mammal Pred Encounter Rate*Stage + Site |
| *Food + Site + Stage* | Probability of use as a function of Food Availability | use ~ Grasshopper Biomass + Pitfall Insect Biomass<br>+ Site + Stage |
| *Food x Stage + Site* | Probability of use as a function of Food Availability depends on Stage | use ~ Grasshopper Biomass*Stage<br>+ Pitfall Insect Biomass*Stage + Site |
| *Vegetation + Site + Stage* | Probability of use as a function of Vegetation Structure | use ~ Bare Ground + Groundcover Height<br>+ Shrub Density + Site + Stage |
| *Vegetation x Stage + Site* | Probability of use as a function of Vegetation Structure depends on Stage | use ~ Bare Ground*Stage + Groundcover Height*Stage<br>+ Shrub Density*Stage + Site |
| *Vegetation<sup>2</sup> + Site + Stage* | Probability of use as a Quadratic function of Vegetation Structure | use ~ Bare Ground + Bare Ground<sup>2</sup><br>+ Groundcover Height + Groundcover Height<sup>2</sup><br>+ Shrub Density + Shrub Density<sup>2</sup> + Site + Stage |
| *Vegetation<sup>2</sup> x Stage + Site* | Probability of use as a Quadratic function of Vegetation Structure depends on Stage | use ~ Bare Ground*Stage + Bare Ground<sup>2</sup>*Stage<br>+ Groundcover Height*Stage + Groundcover Height<sup>2</sup>*Stage<br>+ Shrub Density*Stage + Shrub Density<sup>2</sup>*Stage + Site |
| *Global* | Probability of use as a function of all covariates depends on Stage | use ~ Avian Pred Encounter Rate*Stage<br>+ Mammal Pred Encounter Rate*Stage<br>+ Grasshopper Biomass*Stage<br>+ Pitfall Insect Biomass*Stage<br>+ Bare Ground*Stage + Bare Ground<sup>2</sup>*Stage<br>+ Groundcover Height*Stage<br>+ Groundcover Height<sup>2</sup>*Stage<br>+ Shrub Density*Stage + Shrub Density<sup>2</sup>*Stage<br>+ Site |

The top model output was then examined graphically to better visualize results, alongside violin plots and highest density regions of the raw data.

## RESULTS

### Data summary

At CHBR, we monitored 19 nests (2021, n = 14; 2022, n = 5), and only four broods (2021), due to complete nest failure in 2022. At SP, we monitored 72 nests (2021, n = 38; 2022, n = 34), and 42 broods (2021, n = 21; 2022, n = 21). Locations that were outside of the covariate sampling grid and any broods with less than five locations were censored from analyses (number of brood locations ranged 5-130, median 17). Altogether, our dataset included 83 nest locations (CHBR, n = 19; SP, n = 64) and 39 kernel-density weighted averages for broods (CHBR, n = 3; SP, n = 36), with 122 used units (use = 1) and 1808 available units (use = 0) to fit models.

### Model comparison results

According to our model comparison for this dataset, the most parsimonious model explaining the probability of use by plovers relative to availability was the “Vegetation + Stage + Site” model, carrying 59% of the weight in this model set (Table 2). The second-best model was the “Vegetation x Stage + Site” model, which included an interaction with Stage, carrying 19% of the weight in the model set but at >2 ΔAICc. The top model specifies that habitat use relative to availability has a linear relationship with vegetation structure, when controlling for site and breeding stage differences. The covariates in the top model with well-estimated effects (not overlapping zero) included a positive linear effect of groundcover height, as well as a positive effect of breeding stage (Brood) and a negative effect of Site (SP; Table 3). Results from the second-best, interactive model were within the 95% confidence of the top model, and all interactive terms overlapped zero.

**Table 2.** Model comparison results. Model comparison table to assess competitiveness of hypotheses about the relative influence of environmental covariates on habitat use of Mountain plovers breeding in Colorado.

| Model Name | k<sup>†</sup> | AICc | DAICc | w<sub>i</sub><sup>‡</sup> | Log likelihood |
| --- | --- | --- | --- | --- | --- |
| **Vegetation + Site + Stage** | 6 | 878.89 | 0.00 | 0.59 | -433.42 |
| **Vegetation \* Stage + Site** | 9 | 881.19 | 2.30 | 0.19 | -431.55 |
| **Vegetation<sup>2</sup> \* Stage + Site** | 15 | 881.86 | 2.97 | 0.13 | -425.81 |
| **Vegetation<sup>2</sup> + Site + Stage** | 9 | 883.45 | 4.56 | 0.06 | -432.68 |
| **Global [All covariates]** | 23 | 886.43 | 7.55 | 0.01 | -419.93 |
| **Risk + Site + Stage** | 5 | 887.06 | 8.17 | 0.01 | -438.52 |
| **Food + Site + Stage** | 5 | 890.24 | 11.35 | 0 | -440.10 |
| **Risk \* Stage + Site** | 7 | 890.39 | 11.50 | 0 | -438.16 |
| **Food \* Stage + Site** | 7 | 890.90 | 12.01 | 0 | -438.42 |
| **Intercept** | 1 | 911.87 | 32.98 | 0 | -454.93 |

<sup>†</sup> Number of parameters
<sup>‡</sup> Proportional AICc model weight

**Table 3.** Coefficient values for the top model. Coefficient estimates, standard errors (SE), and 95% confidence intervals (CI) for the top model (*Vegetation + Stage + Site*) from the RSF model selection. Coefficient estimates are on the logit scale. Terms in boldface indicate coefficient estimates with confidence intervals not overlapping zero.

| Parameter | Coefficient | SE | 95% CI | p-value |
| --- | --- | --- | --- | --- |
| (Intercept) | 3.27 | 0.25 | (2.80, 3.75) | <2e-16 |
| Proportion Bare Ground | 0.04 | 0.10 | (-0.14, 0.21) | 0.69 |
| Groundcover Height | **0.40** | **0.12** | **(0.20, 0.61)** | **<0.001** |
| Shrub Density | 0.02 | 0.11 | (-0.17, 0.22) | 0.84 |
| Stage:Brood | **0.76** | **0.20** | **(0.37, 1.16)** | **<0.001** |
| Site:SP | **-1.14** | **0.28** | **(-1.63, -0.65)** | **3.53e-05** |

Specifically, according to the top model, the probability of use for both nest and brood sites increased as height of groundcover vegetation increased (Figure 3a), while no biologically significant effect was detected for proportion bare ground or shrub density (Figure 3b-c). This is contrary to what would be expected given numerous previous studies that suggest plovers use areas with short vegetation and high bare ground during the nesting phase.

**Figure 3.**
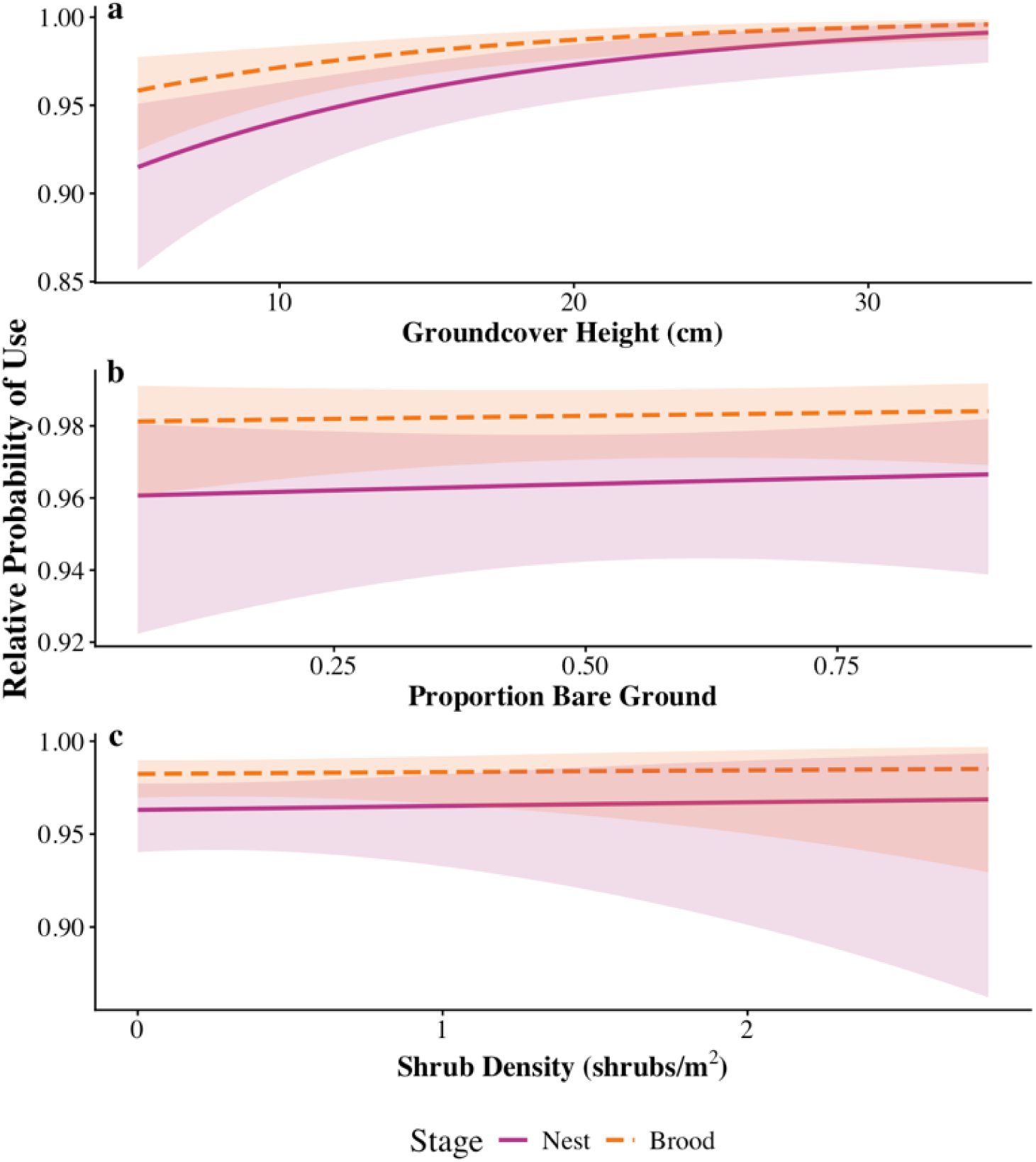
Relative probability of use for nests and broods. Probability of use relative to availability by Mountain plover nests (pink) and broods (orange), as predicted by the top model hypothesizing that habitat use is driven by a linear relationship with vegetation structure. Covariates on the x-axes: (a) Groundcover height (cm), (b) Proportion bare ground, and (c) Shrub density (shrubs per m^2^) are on the original mean-centered scale. Intervals represent 95% confidence.

We examined the used and available sample distributions using violin plots (Figure 4; Figures S6-7) and highest density regions (HDR) grouped by Stage and Site (Figure 5). Both figures suggest that patterns of use differ between the two study sites, but overlap between breeding stages for the most part. Specifically, plovers at CHBR consistently use habitat with shorter vegetation at the lower end of the third quantile and higher bare ground at the upper end of the first quantile of the availability distribution. Meanwhile plovers at SP appear to use most of the availability space during both stages, but with potentially two strategies during the brood-rearing stage: while most broods overlap with the highest density regions of the availability distribution, a smaller sample of broods (n = 5 out of 36) follow a similar pattern to the broods at CHBR.

**Figure 4.**
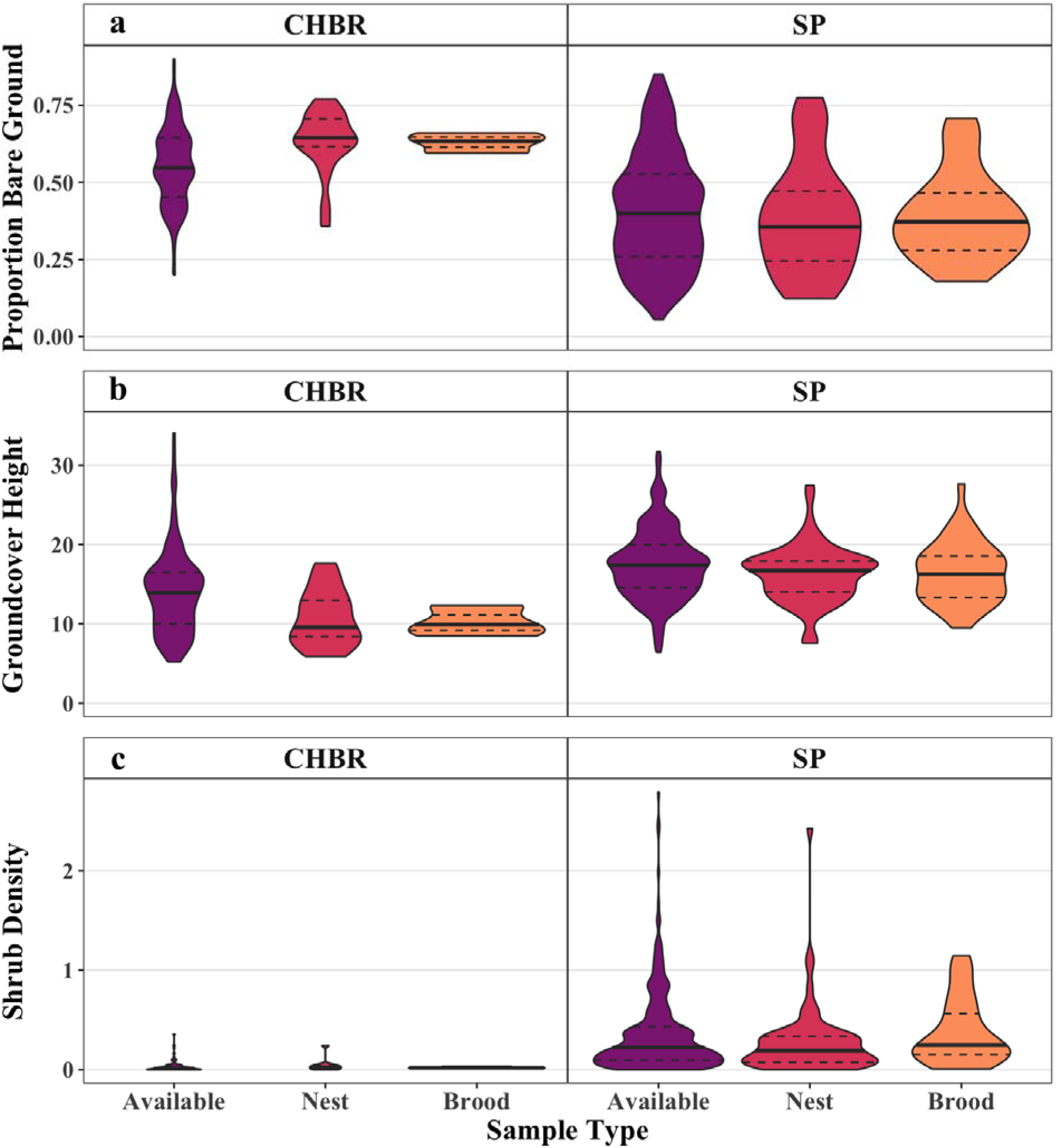
Visualization of vegetation structure at Used vs. Available samples. Violin plots displaying the distributions of the availability samples (purple) alongside samples used by plovers for nests (pink) and broods (orange), for (a) groundcover height (cm), (b) proportion bare ground (0-1), and (c) shrub density (shrubs per m^2^), faceted by Site to show site differences. Bold and dashed lines indicate median and 25% and 75% quantiles, respectively.

**Figure 5.**
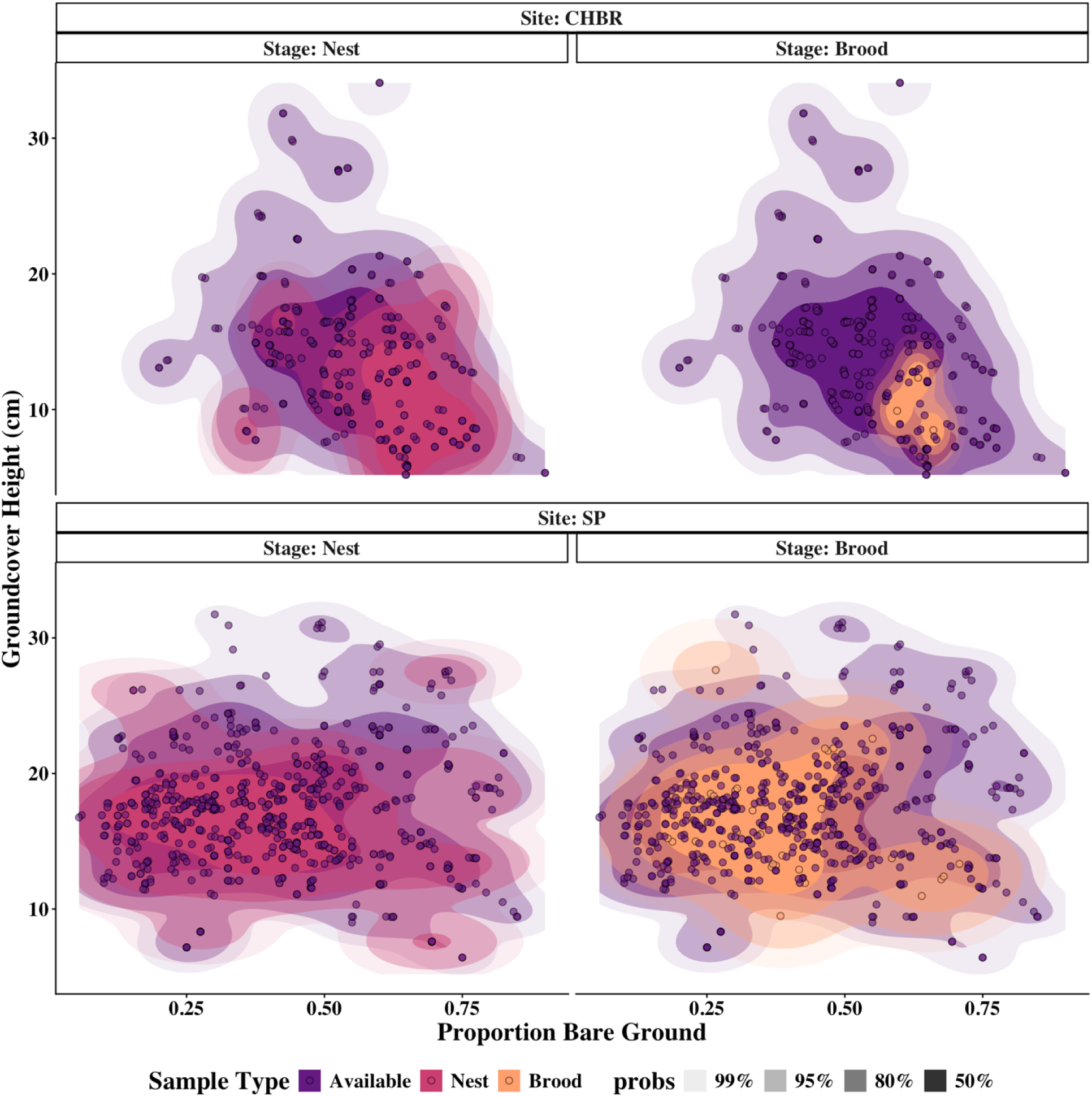
Highest density regions of covariate space for used and available samples. The highest density regions of covariate space (darkest to lightest envelopes: 50, 80, 95, 99% density of sample units) illustrating the relationship between proportion bare ground and groundcover height for the availability distribution (use = 0; purple) and the Used (use = 1) locations of nests (pink) and KD-averaged broods (orange).

The discrepancy between the model output and the patterns in the data is likely a result of the small sample size at CHBR, with the patterns of use of plovers at SP overwhelming those seen at CHBR.

## DISCUSSION

Our study evaluated 10 *a priori* working hypotheses (Table 1) for which of them best explains habitat use relative to availability by the breeding Mountain plover, while investigating the strength of evidence for a dependence on breeding stage. We originally expected that habitat use for breeding Mountain plovers would be best explained by a combination of risk, food, and vegetation that would also depend on breeding stage, thus suggesting stage-specific tradeoffs between risk and rewards. Our model comparison, however, showed that habitat use is best explained by a linear response to spatial variation in vegetation structure while controlling for site and stage differences.

Before we discuss the implications of these results, we acknowledge that the coarse spatial resolution and lack of temporal variation of our covariate indices leave room for alternative explanations. Specifically, the model comparison for this dataset detected no model support for an effect of food availability or predation risk. This should not however be interpreted as plovers not using areas with more food or less risk, as the probability of use is proportional to what is available, and our availability samples were collected only once during the brood-rearing period and at a spatial scale reflecting average daily movement of broods. It is possible that plovers use habitat in response to food, risk, or even vegetation structure, at a finer spatial or temporal scale than we sampled, thus masking potential preferences for or against these indices. These limitations would be expected to *underestimate* the confidence of parameter estimates, and the lack of support for the food and risk hypotheses should therefore be considered with caution. Despite this, the well-estimated effects of vegetation structure from the most parsimonious model provide biologically relevant information that suggests the potential for site-specific and stage-specific shifts in vegetation preferences in the breeding Mountain plover.

The coefficient estimates from the top model, which carried 59% of the weight in our model comparison, suggested a positive relationship with groundcover height during both breeding stages (Figure 3a), which contradicts previous studies of nest site selection that have shown a preference for more bare ground coverage and shorter vegetation (Augustine & Derner, 2012; Duchardt et al., 2020; Knopf & Miller, 1994; Knopf & Wunder, 2023; Manning & White, 2001). However, violin plots of the used and available vegetation samples (Figure 4) as well as the highest density regions of groundcover height and proportion bare ground reveal nuances to the data when stratified by stage and site (Figure 5). Specifically, plovers in CHBR follow the pattern of use found in previous studies, using locations with higher bare ground and shorter vegetation, while plovers in SP appear to use most of the availability space, with a subset of SP broods following a secondary pattern of use like plovers at CHBR. Our models may not have captured these nuances in the data if the limited sample sizes at CHBR being overwhelmed by the much larger sample sizes at SP resulted in a smoothing of the patterns of use.

Despite the limitations of our data, the model comparison indicates that vegetation structure best explains habitat use by plovers during the breeding season, and that a biologically relevant amount of variation is explained by both Stage and Site. There are a few ecological explanations for why plovers might use open vegetation structure (e.g. shorter vegetation, more bare ground) in some contexts but more dense vegetation (e.g. taller vegetation, less bare ground) in others.

Consistent use of locations with more open vegetation structure, as seen at CHBR, might be related to predator vigilance, even though there was little support for an effect of perceived risk. When it comes to nests, selection for specific vegetation features to minimize risk is well-documented in ground-nesting species (Colwell et al., 2005; Hood & Dinsmore, 2007; Knopf & Miller, 1994; Skrade & Dinsmore, 2013; Winton et al., 2000; Webber et al., 2013). Mountain plovers spend 4-14% of the incubation phase off the nest (Skrade & Dinsmore, 2012, but see Graul, 1975), and parental activity such as leaving the nest increases risk of nest predation (Martin et al., 2000). Visibility of the surrounding area might thus be of high importance for plovers when selecting nest sites, as has been demonstrated in the Piping plover (Dorsey et al., 2025). These works, together with the body of previous studies on Mountain plover nest site selection, suggest that the plover might be selecting nest sites where incubating adults can easily anticipate the approach of a predator from hundreds of meters away, regardless of the overall perceived risk landscape.

The very small sample size of broods using open vegetation limits our ability to make inferences about this strategy for broods. However, in a prairie-dog dominated habitat similar to CHBR, Sordahl (1991) found that brood capture sites in northeastern Colorado have higher bare ground coverage and lower vegetation height than control sites, complementing the patterns seen in our own dataset for CHBR (Figure 4; Figure 5, top panels). It may be advantageous to use “open” habitat at this site, and others like it, because they are occupied and maintained by prairie dogs. The highly vigilant prairie dog broadcasts vocal information about risk in real time, from which species like the cooccurring Burrowing owl adjust their own vigilance behaviors (Bryan & Wunder, 2014). Although we did not measure prairie-dog density at CHBR, they were present across most of the site. We propose that at sites like this, plover broods might use the open, shorter vegetation on or near prairie-dog colonies so they can see incoming predators and also eavesdrop on sentinels, as seen with the owl.

At SP, meanwhile, the positive effect of groundcover height suggested by the top model complements what is seen in the violin plots and HDR figures for this site (Figure 4; Figure 5, bottom panels), which show that both nesting and brood-rearing plovers in our SP dataset overlap most of the availability distribution. Our results both complement and expand on a previous study at the same site, in which plovers with broods selected lower bare ground coverage than pre-nesting and post-nesting adults without chicks (Schneider et al., 2006). Yet the highest density regions for what broods use also suggest that, while the majority use “dense” habitat with medium vegetation height and less than 50% bare ground, some broods may still be using areas with high bare ground coverage and short vegetation, as seen at sites like CHBR and Pawnee National Grasslands (Sordahl, 1991; Figure 5). We hypothesize that this more certainly estimated “dense” vegetation strategy seen at SP may be attributed to predation avoidance.

While the hypothesis that predation risk best explains habitat use had little strength of evidence in our model comparison, this could reflect that 1) habitat use does not depend on perceived predation risk, 2) plovers may be unable to assess risk in real time, as it shifts in space and time, or 3) that our risk indices were not fine-scale enough to capture the response of plovers to these covariates. One possibility is that, rather than use areas with lower risk, plovers with chicks may instead be using “dense” vegetation to avoid predation when an encounter occurs. The use of certain vegetation features to minimize risk during incubation or brood-rearing has been well documented in numerous ground-nesting species (Colwell et al., 2005; Hood & Dinsmore, 2007; Knopf & Miller, 1994; Skrade & Dinsmore, 2013; Winton et al., 2000; Webber et al., 2013). Younger Mountain plover chicks also display a hiding strategy when located in denser vegetation during experimental “predator” encounters (Sordahl, 1991), while Schneider et al. (2006) showed that plovers with broods at SP prefer less bare ground and more shrub coverage compared to plovers without broods. In addition, the milder weather at SP diminishes the need of sun-sheltering vegetation at this site., and there was no model support for an effect of food availability in our analysis. These previous and current results altogether support the proposal that this species may be using taller and senser vegetation at SP to camouflage incubating adults as well as young during unpredictable predator encounters, while a smaller number of broods may employ an “open” vegetation strategy that enables adults to anticipate predator encounters from farther away.

## CONCLUSION

Overall, the results from our analysis suggest that the Mountain plover places the most importance on the density of the vegetation structure during the breeding season. While our small sample size, especially at CHBR, limits our ability to make inferences about site- and stage-specific interactive effects on use, an examination of the distributions of used and available samples suggests that vegetation preferences differ between sites and possibly during the brood-rearing stage at SP, although both proposals will require more data to evaluate. Behavioral studies assessing the response of this species to prairie-dog alarm calls would also bring insight to the apparent association where the species cooccur (Augustine & Derner, 2012; Duchardt et al., 2020; Tipton et al., 2009). Our study highlights variable habitat use patterns between geographically separated breeding populations and during the critical brood-rearing phase. We emphasize the importance of considering site-specific and stage-specific habitat requirements when designing management strategies for declining species.

## Funding Statement

Funding was awarded by Colorado Field Ornithologists, Denver Audubon’s Lois Webster Fund, Association of Field Ornithologists, and Bird Conservancy of the Rockies. Colorado State University – Pueblo awarded SEED grant funding to CWVR. Colorado Parks & Wildlife provided camera trap equipment.

## Acknowledgements

We acknowledge the Ranchlands LLC team and Phillips family, Colorado State Land Board, Bureau of Land Management, and Colorado Parks & Wildlife for assisting with access to study sites. We thank field assistants Chrissy Gatian, Shawn Overby, Elissa Velasquez, Jake Powers, and Zoe Erkenbeck. We also thank Jeroen Reneerkens, Camilo Carneiro, and others for taking the time to review this manuscript.

## Conflict of Interest Statement

The authors declare no conflict of interest.

## Supplementary Information

Additional supporting information may be found online in the Supporting Information section at the end of the article.

**Photograph S1. First day of life for a Mountain plover chick.**

**Photograph S2. Nearly fledged.**

**Photograph S3. Last chick of the season. Photograph S4. A worried parent.**

**Figure S1. 2021 Covariate raster for Chico Basin Ranch.**

**Figure S2. 2022 Covariate raster for Chico Basin Ranch.**

**Figure S3. 2021 Covariate raster for South Park.**

**Figure S4. 2022 Covariate raster for South Park.**

**Figure S5. Covariate correlation test results.**

**Figure S6. Visualization of predation risk at Used vs. Available Samples.**

**Figure S7. Visualization of food availability at Used vs. Available Samples.**

**Figure S8. Site-specific relative probability of use as predicted by the top model.**

## Notes

https://figshare.com/articles/dataset/Weissburg_et_al_Mountain_Plover_Stage-specific_habitat_use_-_Markdown_and_data_files/25658022

### Competing Interest Statement

The authors have declared no competing interest.

### Summary of Updates

The manuscript and associated code has been revised for resubmission. We have made what we believe are substantial improvements to the clarity of the methods and the structure of the analysis, as well as to the discussion by providing discourse on alternative explanations of the results due to the limitations of covariate sampling resolution.

