## Supplementary Online Material for "Stage-specific habitat use of the Mountain Plover in Colorado, USA"

**Supporting Information**

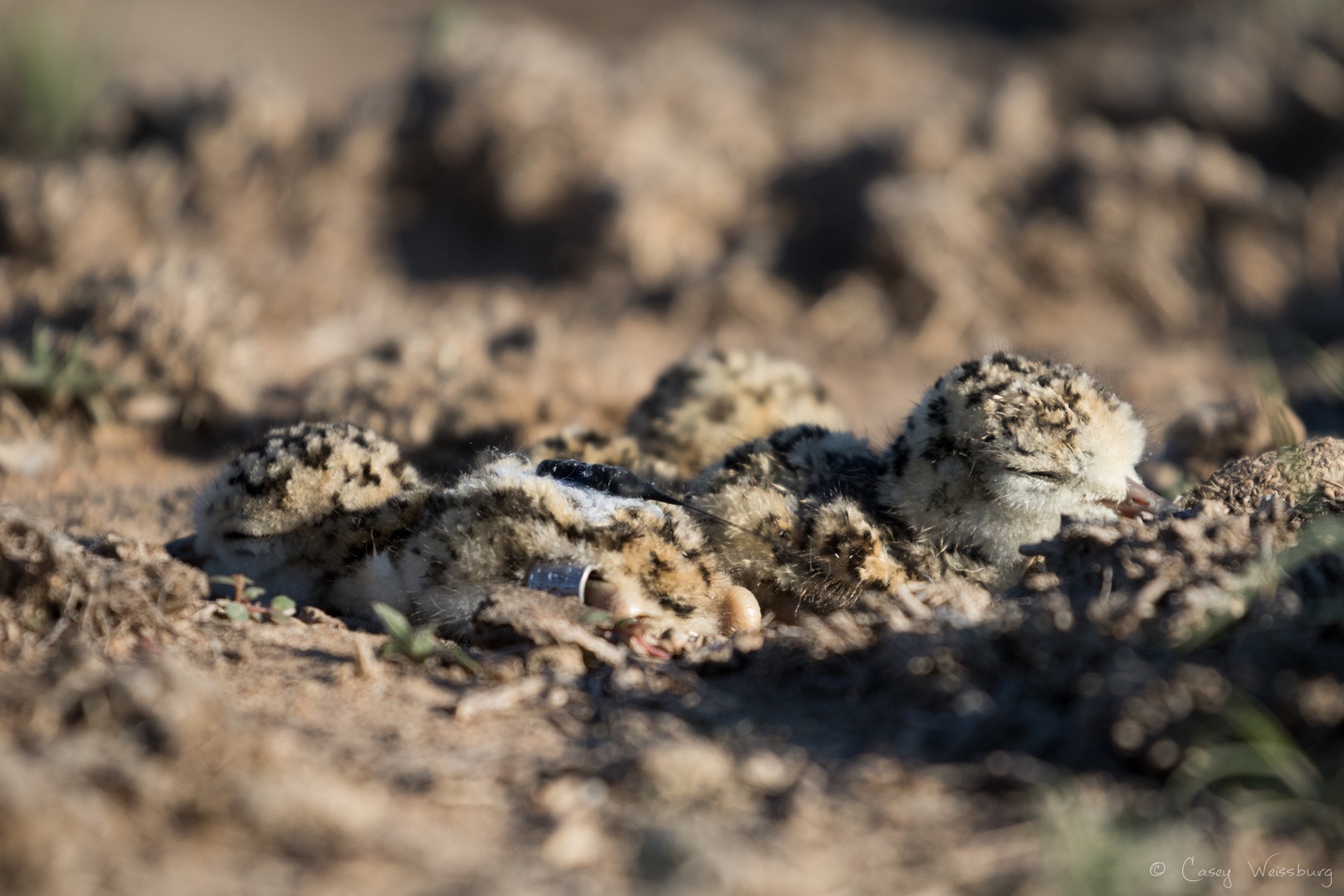

**Photograph S1. First day of life for a Mountain plover chick.** Three Mountain plover chicks, less than one-day old, resting in the nest cup after banding and radio-tagging. Photo by Casey Weissburg.

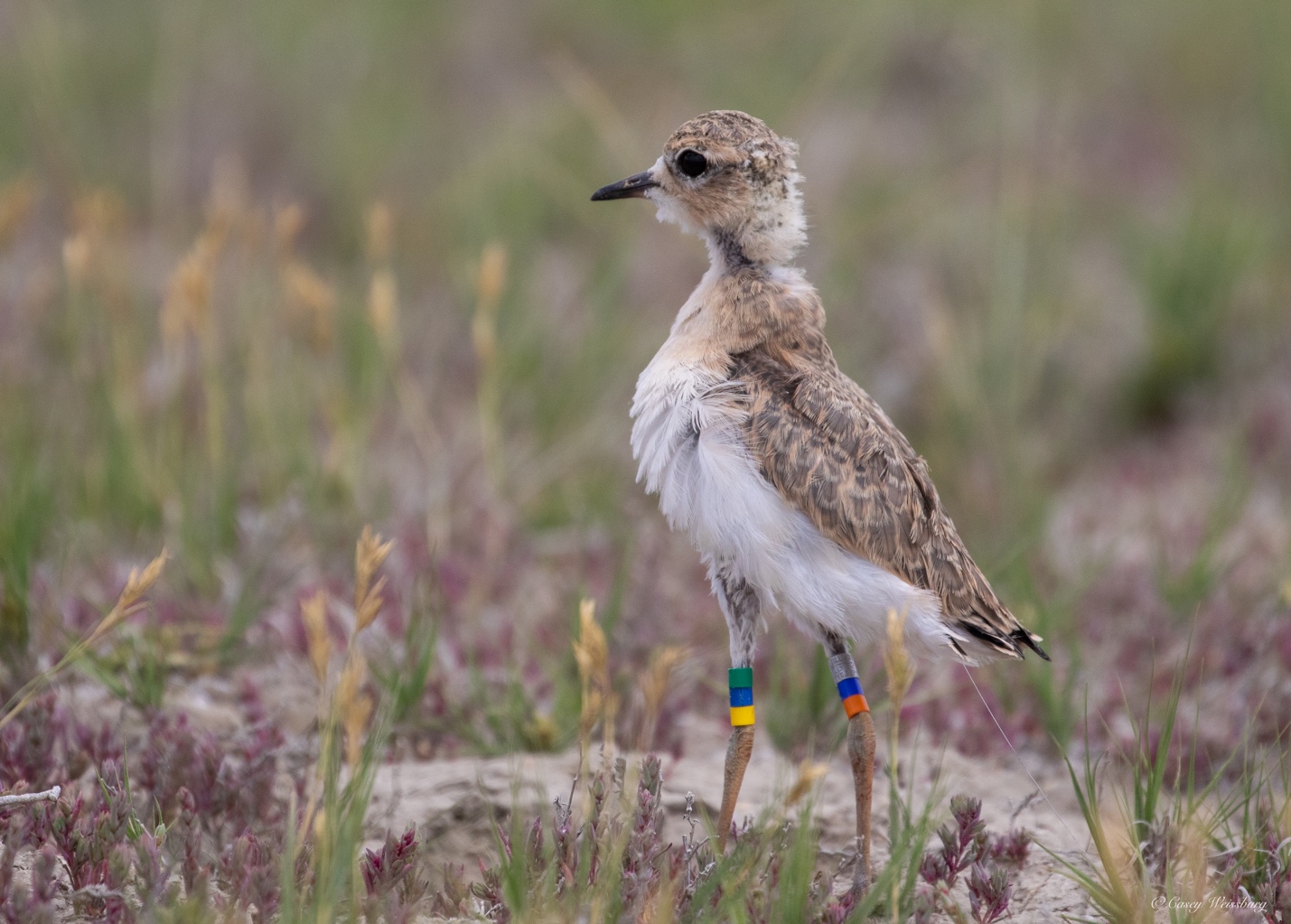
**Photograph S2. Nearly fledged.** A radio-tagged Mountain plover chick walks away after receiving a unique color-band combination in the last week before fledging. Photo by Casey Weissburg.

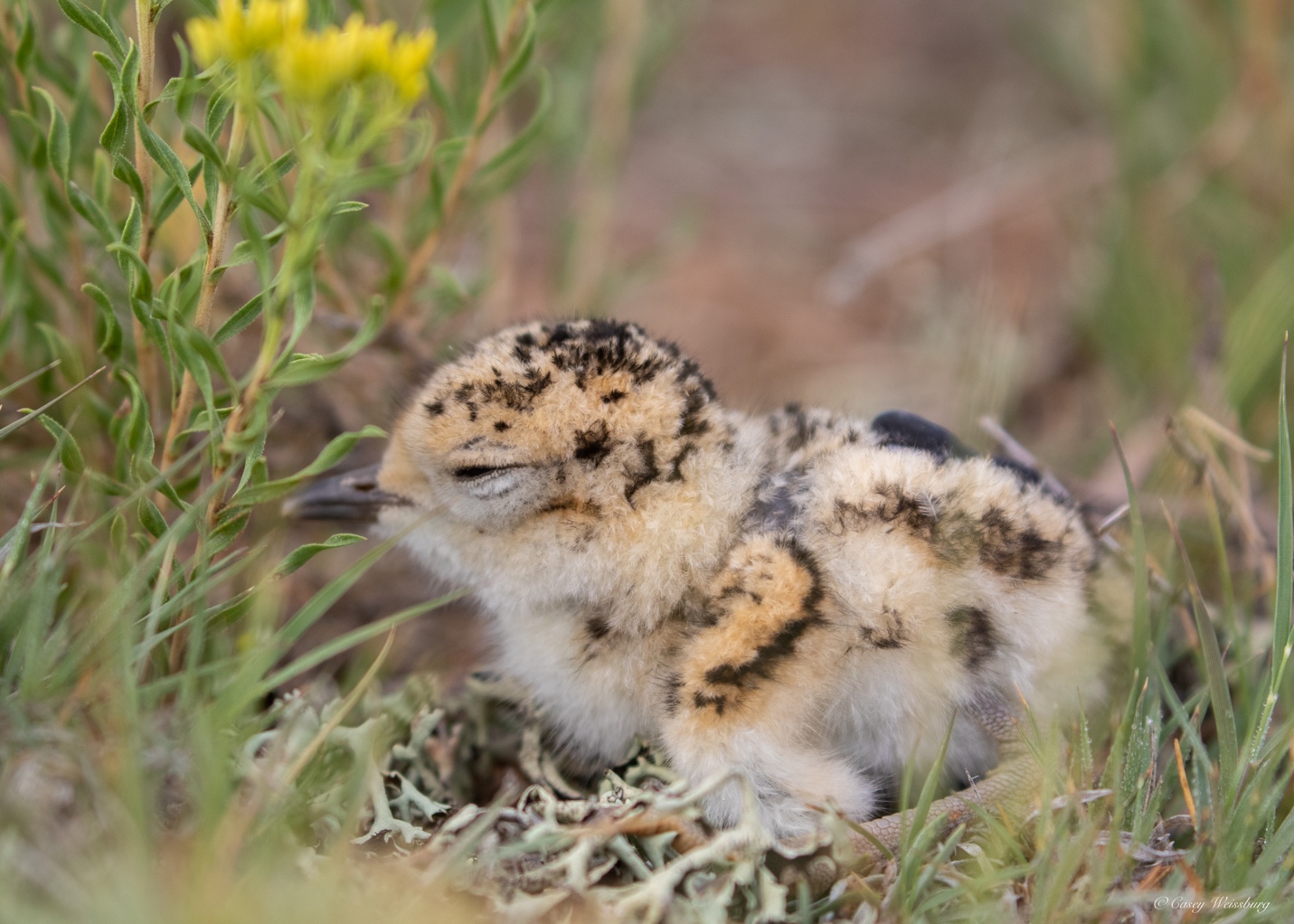
**Photograph S3. Last chick of the season.** A 3-day old chick rests under the rabbitbrush where it was captured, after banding and radio-tagging. Photo by Casey Weissburg.

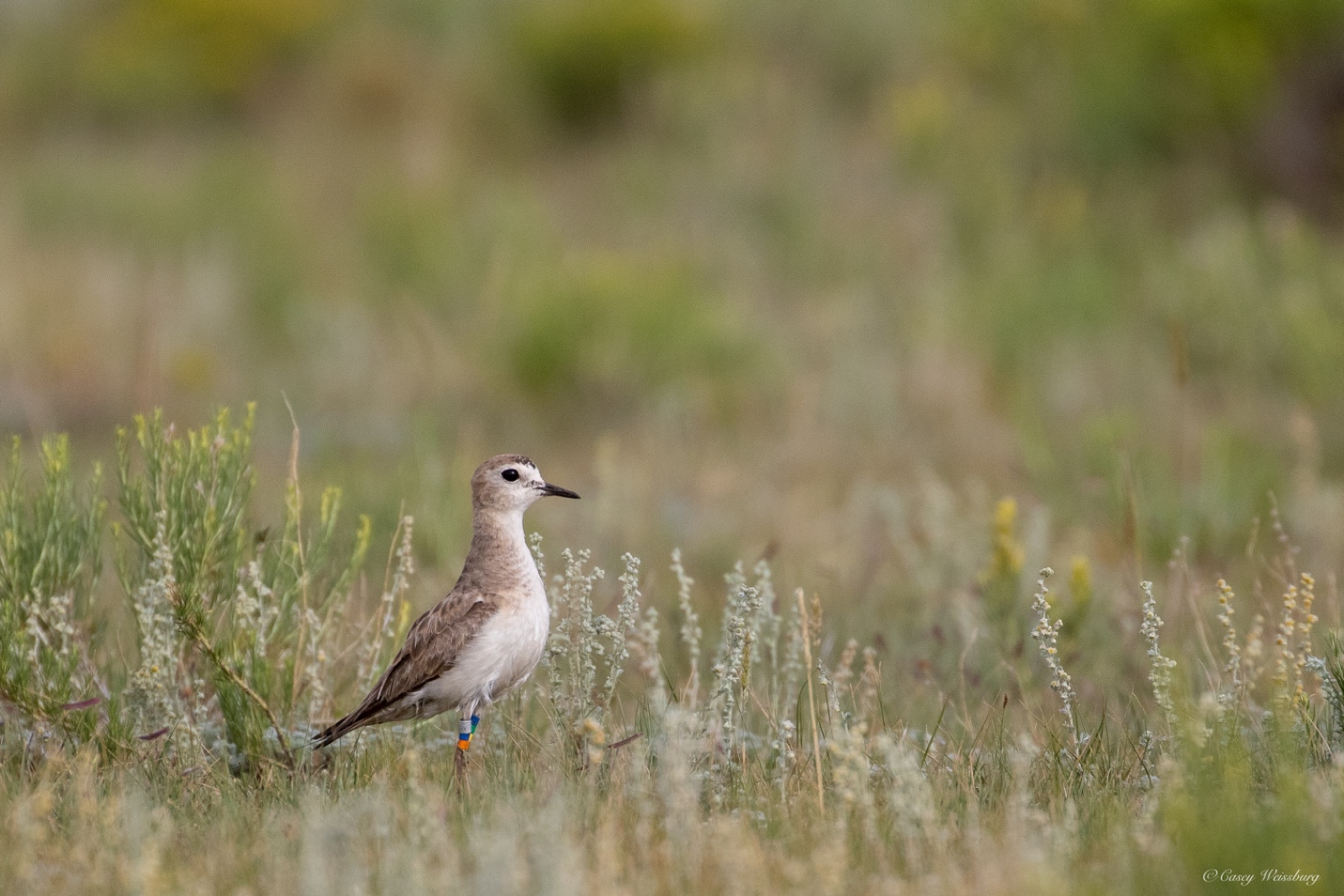
**Photograph S4. A worried parent.** An adult Mountain plover watches from close by while its newly hatched chick is banded and radio-tagged (the same chick from Figure S3). Photo by Casey Weissburg.

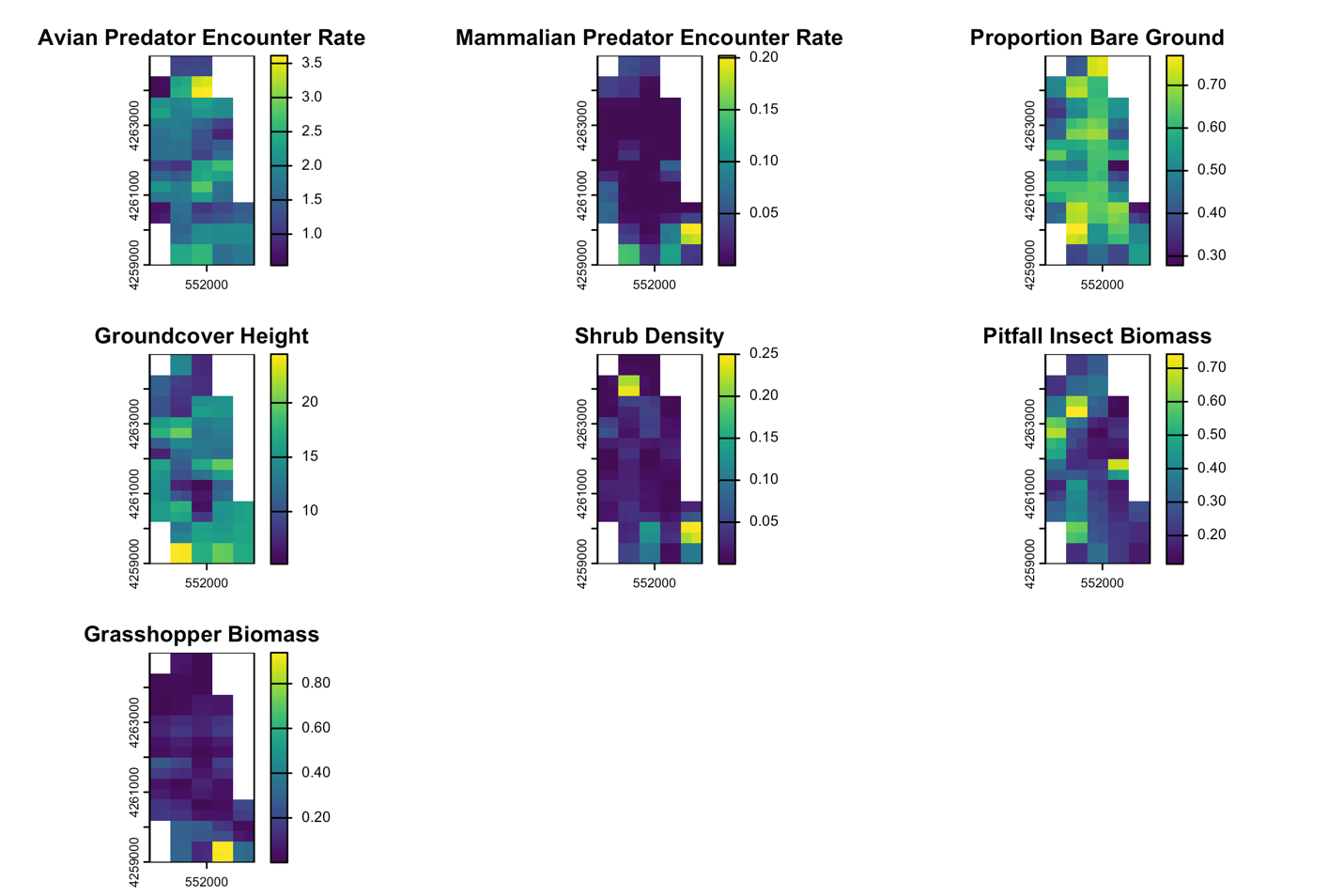

**Figure S1. 2021 Covariate raster for Chico Basin Ranch.** A visualization of environmental covariate data collected at Chico Basin Ranch in 2021, rasterized and estimated to a finer resolution of 300-by-300 meters using inverse distance weighted interpolation in QGIS. Covariates from left to right and top to bottom: Total avian predator encounter rate, Mammalian predator encounter rate, Proportion bare ground coverage, Groundcover height, Shrub density, Pitfall insect biomass, Grasshopper biomass.

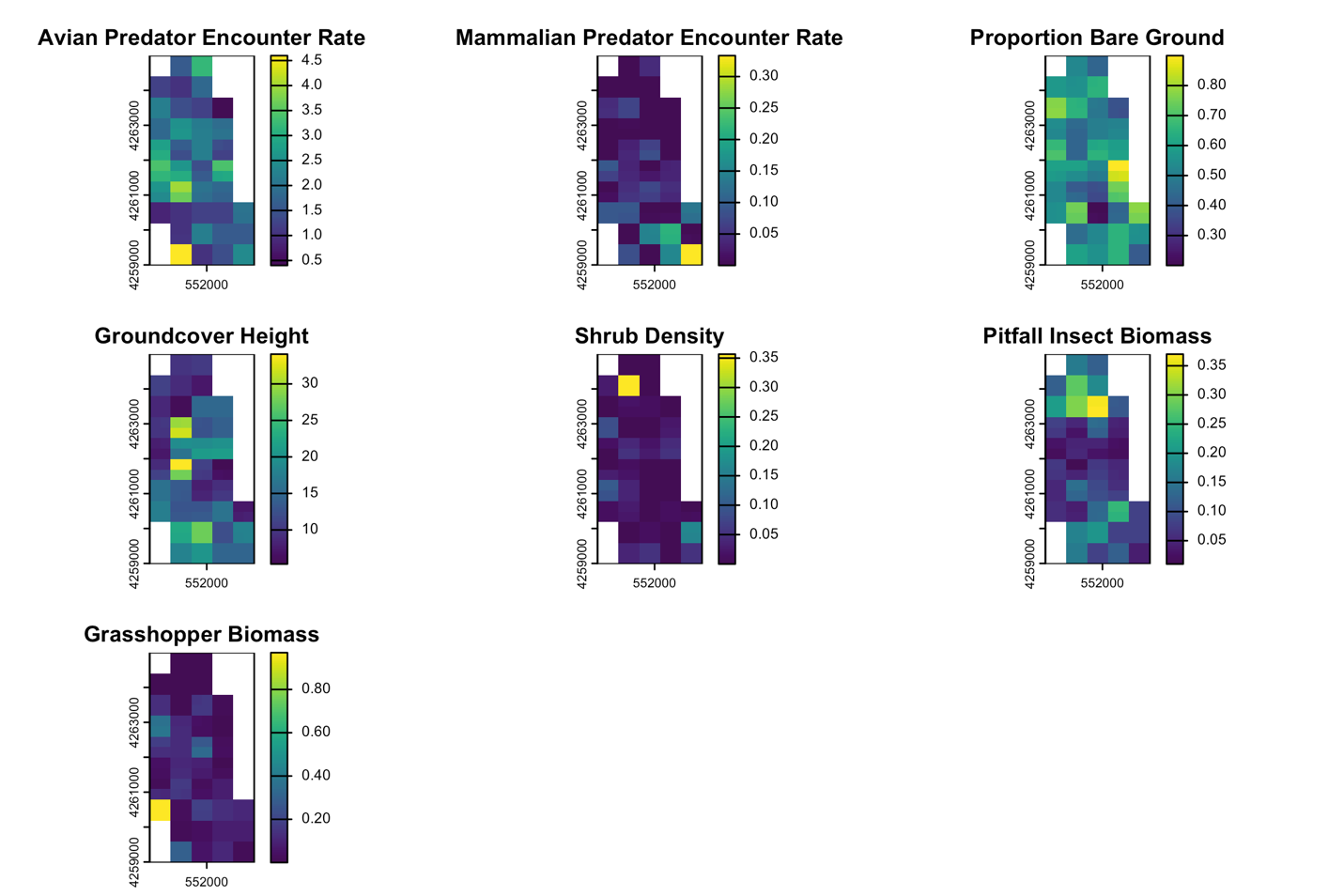

**Figure S2. 2022 Covariate raster for Chico Basin Ranch.** A visualization of environmental covariate data collected at Chico Basin Ranch in 2022, rasterized and estimated to a finer resolution of 300-by-300 meters using inverse distance weighted interpolation in QGIS. Covariates from left to right and top to bottom: Total avian predator encounter rate, Mammalian predator encounter rate, Proportion bare ground coverage, Groundcover height, and Shrub density, Pitfall insect biomass, Grasshopper biomass.

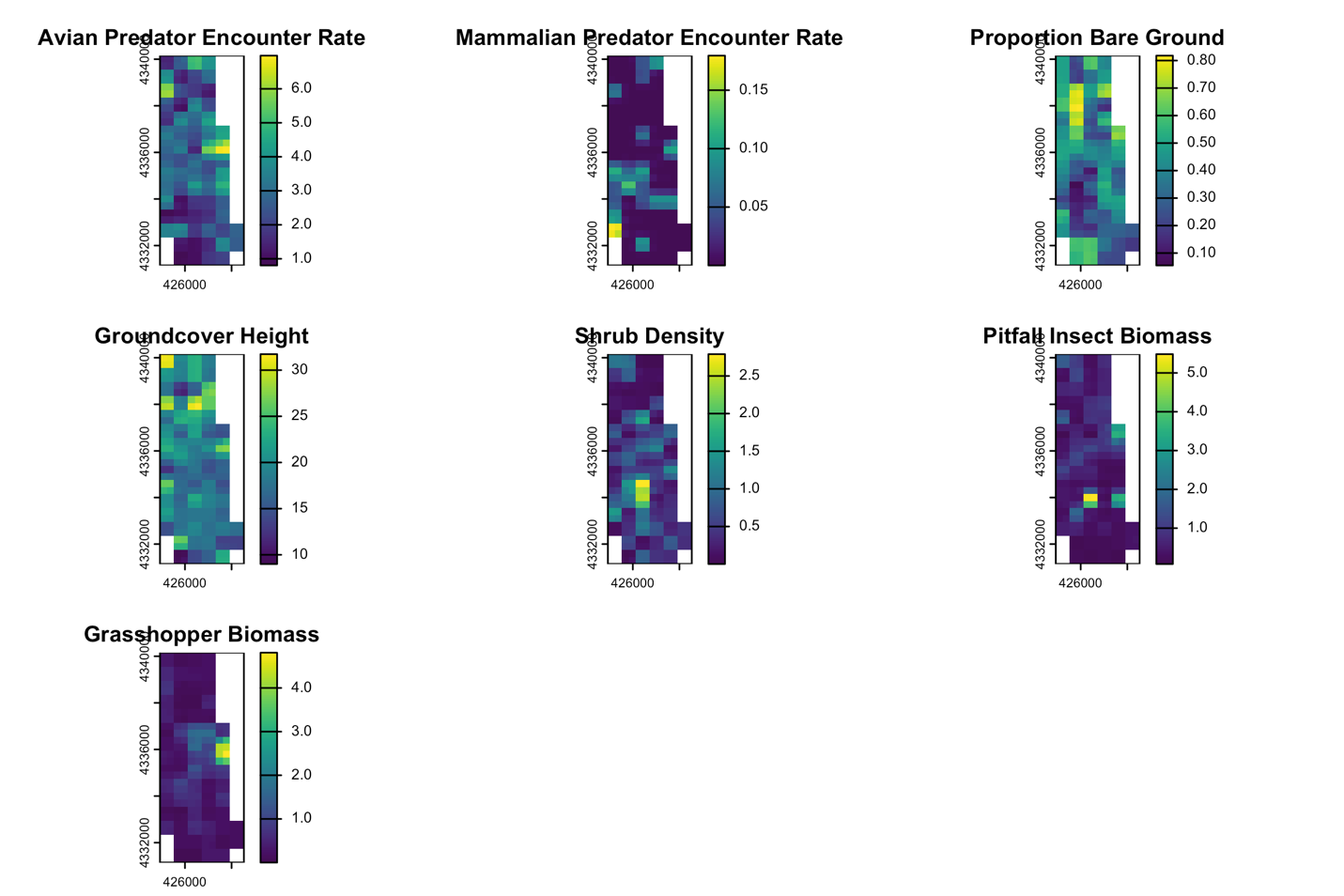
**Figure S3. 2021 Covariate raster for South Park.** A visualization of environmental covariate data collected at South Park in 2021, rasterized and estimated to a finer resolution of 300-by-300 meters using inverse distance weighted interpolation in QGIS. Covariates from left to right and top to bottom: Total avian predator encounter rate, Mammalian predator encounter rate, Proportion bare ground coverage, Groundcover height, Shrub density, Pitfall insect biomass, Grasshopper biomass.

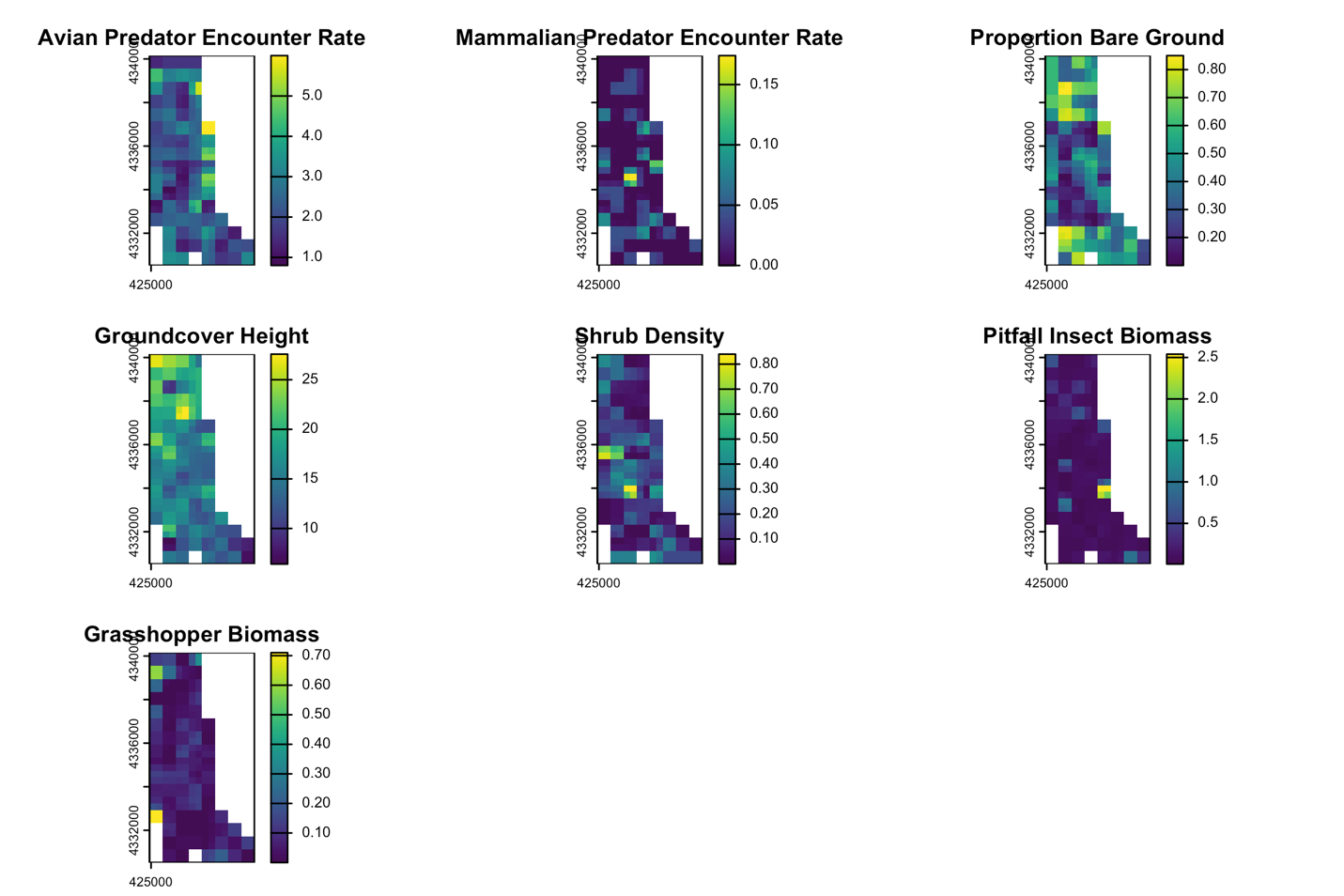

**Figure S4. 2022 Covariate raster for South Park.** A visualization of environmental covariate data collected at South Park in 2022, rasterized and estimated to a finer resolution of 300-by-300 meters using inverse distance weighted interpolation in QGIS. Covariates from left to right and top to bottom: Total avian predator encounter rate, Mammalian predator encounter rate, Proportion bare ground coverage, Groundcover height, Shrub density, Pitfall insect biomass, Grasshopper biomass,.

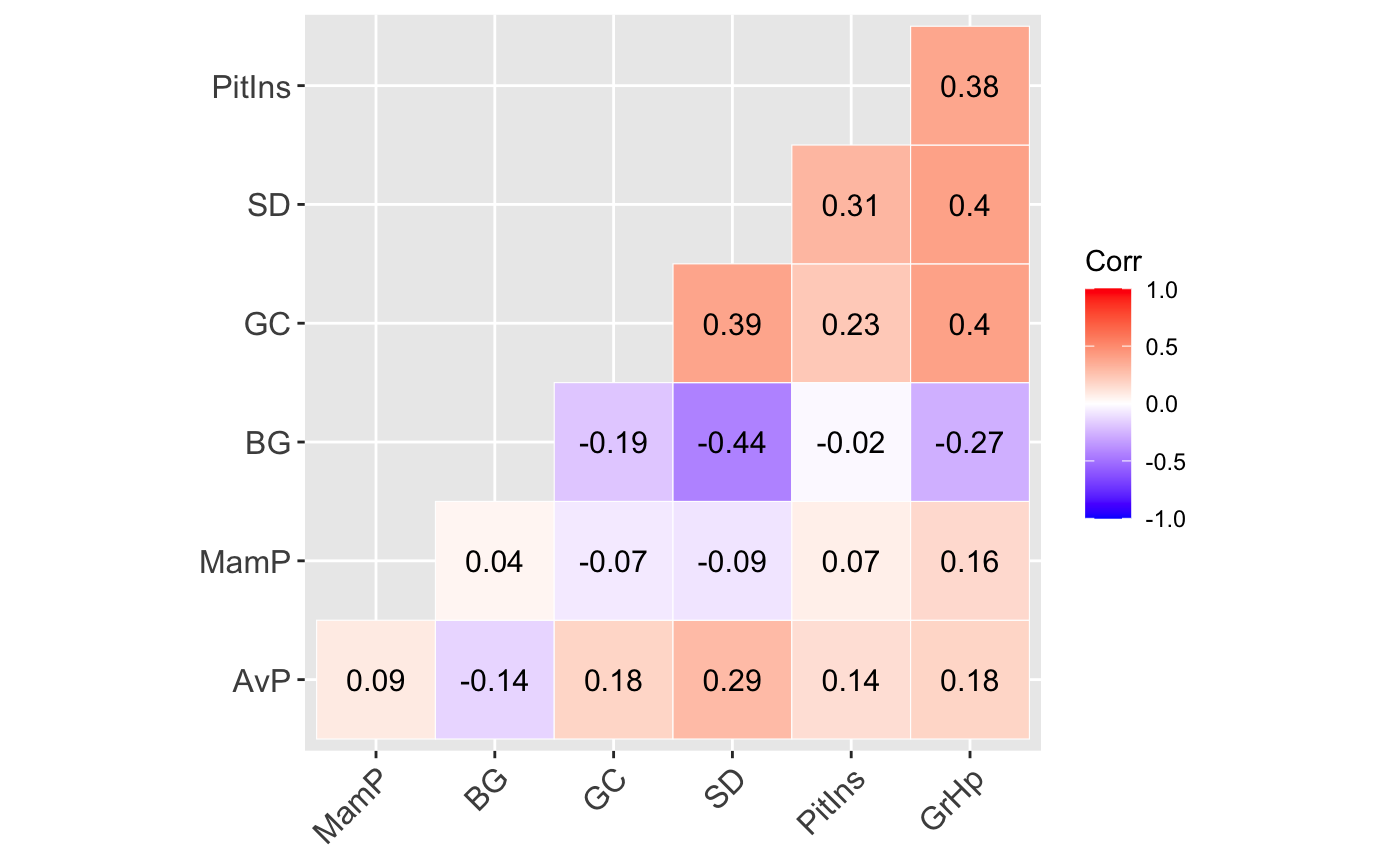
**Figure S5. Covariate correlation test results.** Correlogram of Spearman’s rank correlations (red = negative relationship; blue = positive relationship) between all covariate data for used and available locations. AvP = Avian predator encounter rate; MamP = Mammalian predator encounter rate; BG = Proportion bare ground; GC = Groundcover height; SD = Shrub density; PitIns = Pitfall insect biomass; GrHp = Grasshopper biomass.

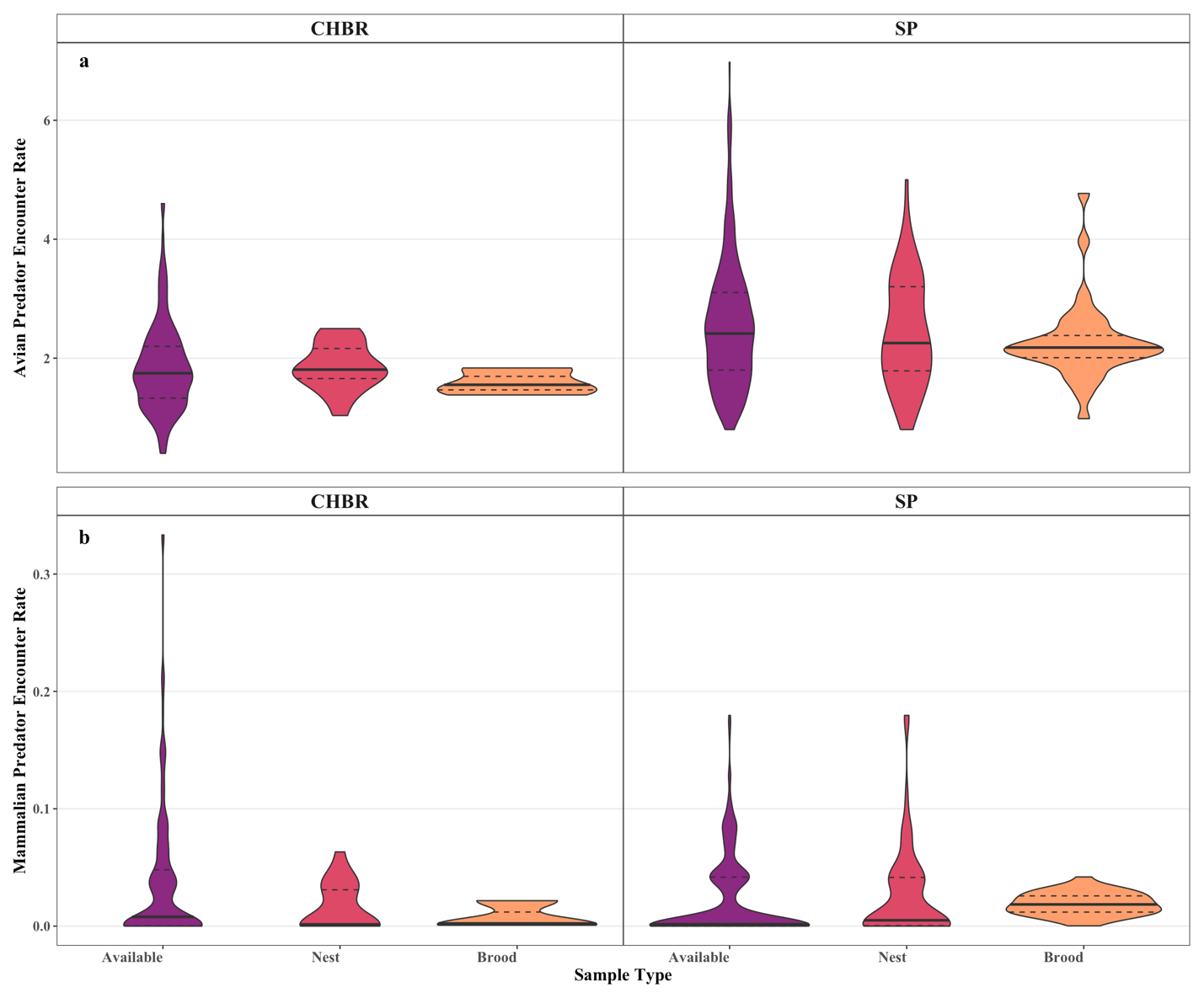

**Figure S6.** **Visualization of predation risk at Used vs. Available samples.** Violin plots displaying the distributions of the availability samples (purple) alongside samples used by plovers for nests (pink) and broods (orange), for (a) avian predator encounter rate (birds per survey), and (b) mammalian predator encounter rate (encounters per active camera night), faceted by Site to show site differences. Bold and dashed lines indicate median and 25% and 75% quantiles, respectively.

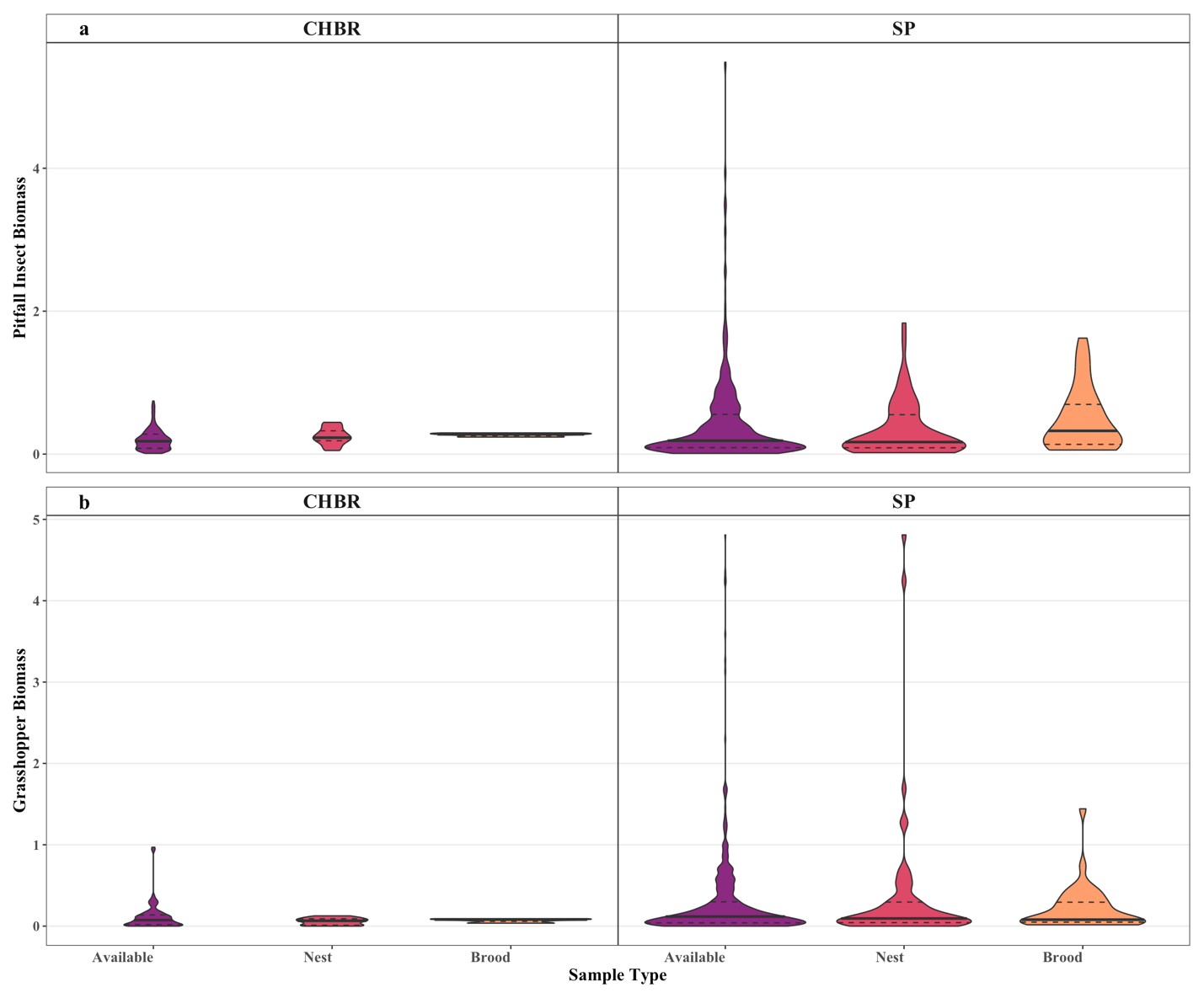

**Figure S7. Visualization of food availability at Used vs. Available samples.** Violin plots displaying the distributions of the availability samples (purple) alongside samples used by plovers for nests (pink) and broods (orange), for (a) pitfall insect biomass (dry grams per effort), and (b) grasshopper biomass (dry grams), faceted by Site to show site differences. Bold and dashed lines indicate median and 25% and 75% quantiles, respectively.

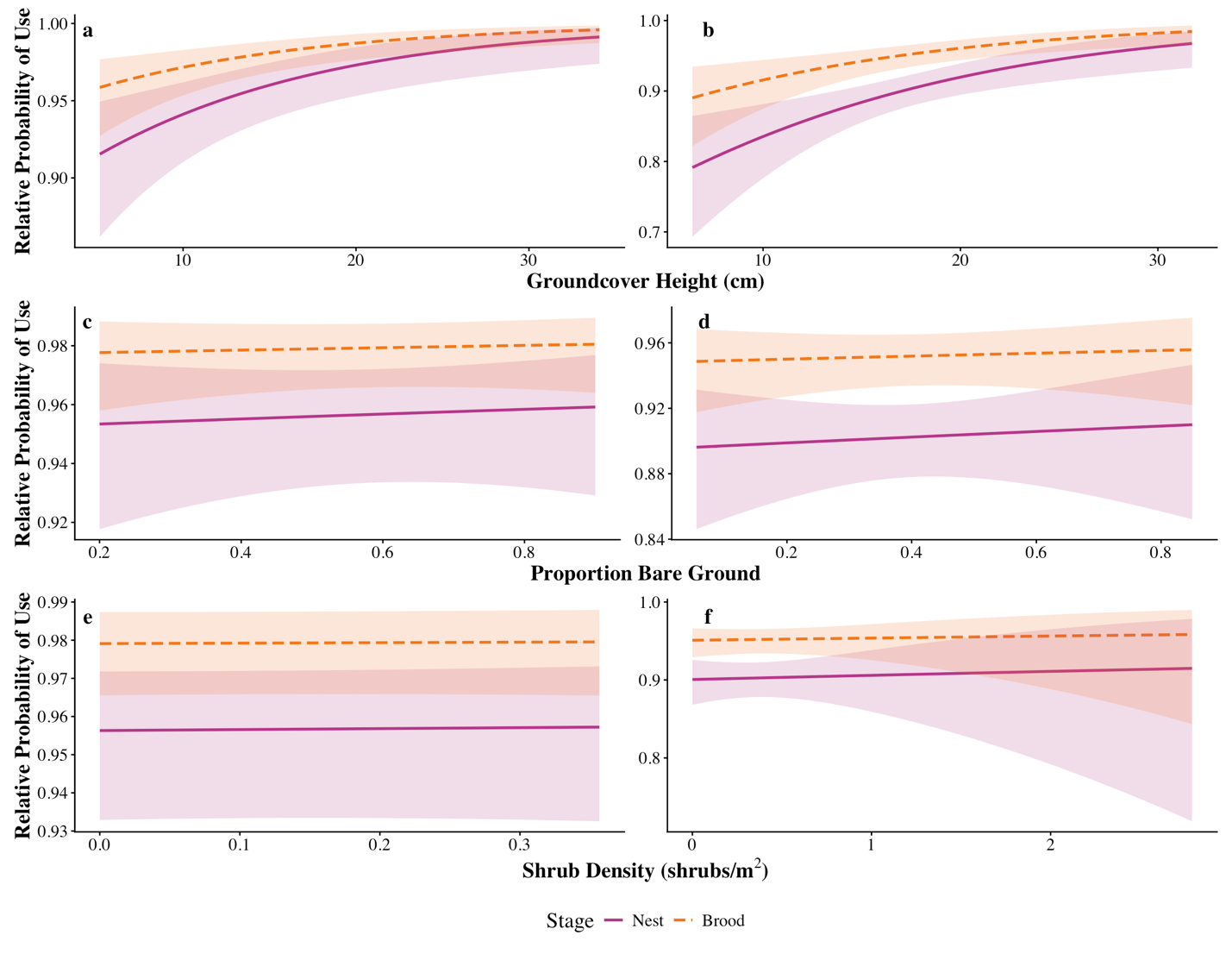

**Figure S8. Site-specific relative probability of use as predicted by the top model.**  Probability of use relative to availability by Mountain plover nests (pink) and broods (orange) at CHBR (left) and SP (right), as predicted by the top model hypothesizing that habitat use is driven by a linear relationship with vegetation structure while conditioning on breeding stage and site. Covariates (a-b) Groundcover height (cm), (c-d) Proportion bare ground, and (e-f) Shrub density (shrubs/m^2^) are on the original scale. Intervals represent 95% confidence.
